# A stochastic model for error correction of kinetochore-microtubule attachments and its coupling to the spindle assembly checkpoint

**DOI:** 10.1101/541573

**Authors:** Anand Banerjee, Neil Adames, Jean Peccoud, John J. Tyson

## Abstract

To divide replicated chromosomes equally between daughter cells kinetochores must attach to microtubules emanating from opposite poles of the mitotic spindle. Two mechanisms, namely, error correction and ‘spindle assembly checkpoint’ work together to facilitate this process. The error correction mechanism recognizes and detaches erroneous kinetochore-microtubule attachments, and the spindle assembly checkpoint delays the onset of anaphase until all the kinetochores are properly attached. Kinases and phosphatases at the kinetochore play a key role in proper functioning of these two mechanisms. Here we present a stochastic model to study how the opposing activities of kinases and phosphatases at the kinetochore affect error correction of kinetochore-microtubule attachments and checkpoint signaling in budding yeast, *Saccharomyces cerevisiae*. We show that error correction and biorientation of chromosomes occurs efficiently when the ratio between kinase activity of Ipl1 and the activity of an opposing phosphatase is a constant (balance point), and derive an approximate analytical formula that defines the balance point. Analysis of the coupling of the spindle assembly checkpoint signal to error correction shows that its strength remains high when the Ipl1 activity is equal to (or larger than) the value specified by the balance point, and the activity of another kinase, Mps1, is much larger (approximately 30 times larger) than its opposing phosphatase (PP1). We also find that the geometrical orientation of sister chromatids does not significantly improve the probability of their reaching biorientation, which depends entirely on Ipl1-dependent microtubule detachment.

**Author summary:** The kinetochore, the master regulator of chromosome segregation, integrates signals from different chromosome attachment states to generate an appropriate response, like the destabilization of erroneous kinetochore-microtubule attachments, stabilization of correct attachments, maintenance of the spindle assembly checkpoint signal until all kinetochores are properly attached, and finally silencing of checkpoint when biorientation is achieved. At a molecular level the job is carried out by kinases and phosphatases. The complexity of the interactions between these kinases and phosphatases makes intuitive analysis of the control network impossible, and a systems-level model is needed to put experimental information together and to generate testable hypotheses. Here we present such a model for the process of error correction and its coupling to the spindle assembly checkpoint in budding yeast. Using the model, we characterize the balance between kinase and phosphatase activities required for removing erroneous attachments and then establishing correct stable attachments between kinetochore and microtubule. We also analyze how the balance affects the strength of the spindle assembly checkpoint signal.

## Introduction

Equal partitioning of duplicated chromosomes is crucial for maintaining genetic integrity from one generation to the next. A key step in this process is the attachment of kinetochores (KTs) on sister chromatids to microtubules (MTs) emanating from opposite poles of the mitotic spindle. The attachment process is stochastic and error prone, resulting in erroneous attachments like syntely (where both KTs are attached to the same spindle pole) and merotely (where one KT is attached to both spindle poles) (see Fig 1). Such errors must be corrected before the onset of anaphase (1–3). The correction of erroneous KT-MT attachments in budding yeast is crucially dependent on the kinase Ipl1 (3–5).

**Fig 1.**
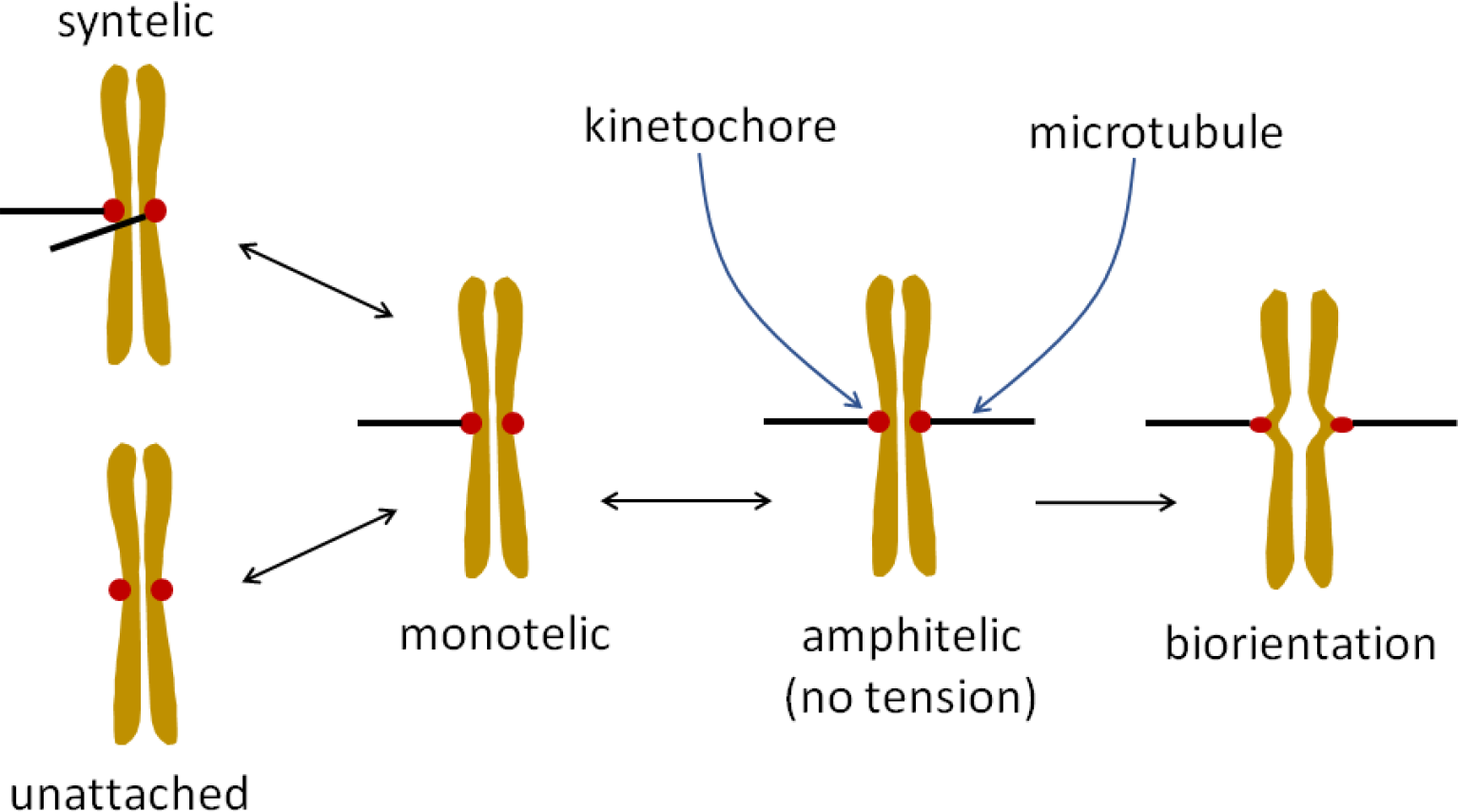
Model for transitions of sister chromatids between different KT-MT attachment states. Double-headed arrows denote reversible transitions, whereas the single-headed arrow denotes the irreversible transition from the amphitelic to the biorientation state.

The Ndc80 complex at the KT is a primary site for KT-MT attachment (6, 7). Phosphorylation of Ndc80 by Ipl1 weakens its interaction with MTs (7–9) and conversely, its dephosphorylation increases its affinity for MTs and stabilizes KT-MT attachments (10). In tensionless KTs (which is the case in unattached, syntelic and monotelic KTs), Ipl1 phosphorylates Ndc80, resulting in dissociation of Ndc80-MT interactions. This provides the unattached KT an opportunity to attach to a new MT from the correct spindle pole (1, 3, 4). Together, these observations suggest that a balance between kinase and phosphatase activities is required to break erroneous attachments and then establish correct, stable attachments between KT and MT (11–13). Experiments show that PP1 phosphatase (Glc7 is the PP1 catalytic subunit in budding yeast) opposes the kinase activity of Ipl1 (14–16), but whether it dephosphorylates Ndc80 or not and its importance in biorientation of chromosomes remains unclear.

The process of error correction is also coupled to a surveillance mechanism called the spindle assembly checkpoint (SAC). Unattached KTs, generated during error correction, initiate SAC signaling (17, 18). Briefly, unattached KTs recruit a kinase, Mps1, to phosphorylate Spc105 (Knl1 in mammalian cells) at phosphodomains called MELTs (Met-Glu-Leu-Thr sequence). Phosphorylated MELTs then bind to cytoplasmic SAC proteins to turn on the SAC signal (19). The SAC monitors KT-MT attachments and delays the onset of anaphase until all KTs are properly attached (17, 20). After biorientation is achieved, the SAC needs to be turned off to allow cells to proceed to anaphase. KTs recruit phosphatases PP1 and PP2A (Pph21/22 are the PP2A catalytic subunits in yeast) to silence the SAC in a timely manner (21–23). Almost all the proteins involved in error correction and the SAC are conserved in yeast and humans. However, yeast KTs are smaller than mammalian KTs and attach to only one MT (24).

The activities of these kinases and phosphatases are coupled to each other. For example, Ipl1 and PP2A activities control the binding of PP1 to KTs, and the attachment of Ndc80 to a MT blocks its Mps1 binding sites (25, 26). Existing models on error correction are coarse-grained and do not take into account the complexity of interactions between the kinases and phosphatases at the KT (27, 28). They also do not account for how SAC signaling is coupled to the error correction process. To fill this gap, in this paper we present (in the context of budding yeast cells) a new systems-level, stochastic model to track the time evolution of the number of molecules of kinases and phosphatases bound to the KT, the phosphorylation states of their substrates, and the number of SAC proteins bound to the KT. We study how the opposing activities of kinases and phosphatase affect error correction in KT-MT attachments and the activity of the SAC. We also calculate the relative contributions of Ipl1-dependent destabilization of KT-MT attachments and the geometrical orientation of KTs towards reaching biorientation.

## Model

The scheme in Fig 1 shows the possible transitions between different attachment states for a pair of budding yeast KTs at the centromere of a replicated chromosome. Initial attachment of a KT to a MT occurs with the lateral surface of the MT – known as lateral attachment. The lateral attachment is then converted into end-on attachment by sliding of the KT on the MT. To simplify this process, we assume that the first interaction between KT and MT is an end-on attachment. In our model, the amphitelic state has the KTs attached to MTs from opposite poles of the mitotic spindle but the centromere is not yet under tension. The amphitelic state can transition reversibly to the monotelic state (only one KT attached to the spindle) or irreversibly to the biorientation state in which the KTs are attached to opposite poles and there is tension between the KTs. Tension is generated by the opposing forces exerted by depolymerizing MTs bound to KTs. Tension stretches the centromeric region of the chromosome and is thought to stabilize KT-MT attachments by physically separating Ipl1 from its substrates and thereby reducing its role in destabilizing KT-MT attachments (29, 30). We assume that Ipl1 activity behaves like a step function: in the amphitelic state, Ipl1 activity remains high, which allows the possibility of detachment of a MT, and in biorientation state it drops to zero. Hence, the probability of going from the biorientation state to the amphitelic state is zero. The criterion used to define the transition from the amphitelic to biorientation state is described later.

The actions of kinases and phosphatases at the KT to control error correction and SAC activity, as described above, is shown schematically in Fig. 2. The main kinases are Ipl1 and Mps1, and the phosphatases are PP1, PP2A, and PPX (an unknown phosphatase). Ipl1 phosphorylates Ndc80 and the RVSF motif on Spc105 (8, 16). Phosphorylation of RVSF prevents its binding to PP1. Mps1 kinase phosphorylates MELT repeats on Spc105 to activate SAC signaling (22). PP2A dephosphorylates the RVSF motif, allowing PP1 to bind to Spc105 and dephosphorylate its MELT repeats. At this point in time, it is not clear which phosphatase dephosphorylates Ndc80; hence, ‘PPX’ in Fig. 2. Possible candidates for PPX are the free PP1 and PP2A in the nucleus.

**Fig 2.**
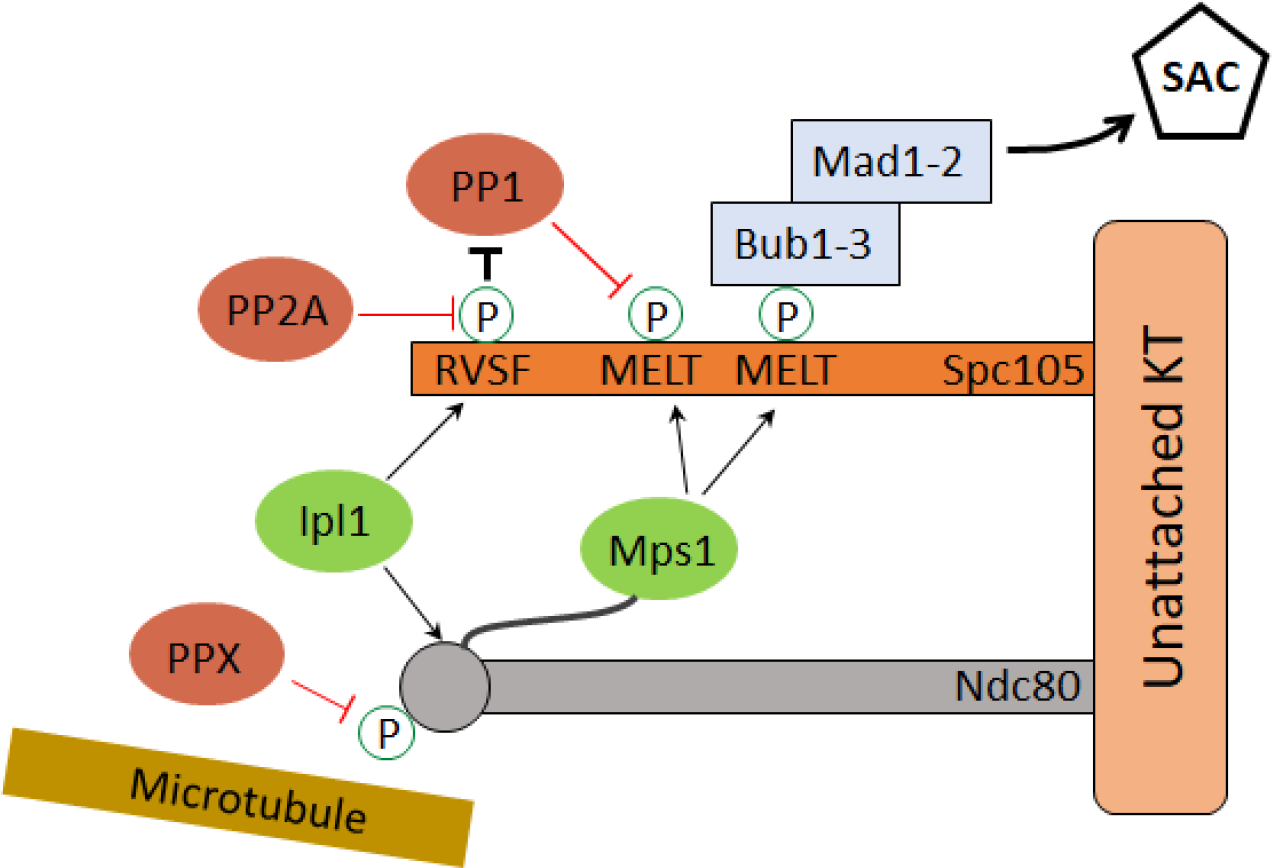
Scheme for kinase and phosphatase activities at the KT. Kinases are Ipl1 and Mps1 (shown in green), and phosphatases are PP1, PP2A, and PPX (shown in red). Phosphorylation of MELT motifs by Mps1 starts the SAC signaling cascade. Phosphorylation of Ndc80 by Ipl1 weakens KT-MT attachment, and phosphorylation of RVSF by Ipl1 prevents binding of PP1. PP2A dephosphorylates RVSF to promote binding of PP1 to Spc105. Dephosphorylation of Ndc80 by PPX promotes KT-MT attachment, and dephosphorylation of MELT motifs by PP1 promotes silencing of the SAC signal.

The scheme shown in Fig. 2 can be understood as three coupled modules, namely, the Ndc80 module, the RVSF module, and the MELT module. These modules and the coupling between them are shown in Fig. 3A. The Ndc80 module consists of two phosphorylation states and two Mps1-binding states. Ipl1 phosphorylates Ndc80 at multiple sites to modulate its interactions with MTs (8); to keep the model simple we assume only two phosphorylation states. Attachment of a MT to Ndc80 is diagrammed separately (Fig. 3B). A budding yeast KT contains five Ndc80 molecules (31). The unknown phosphatase PPX dephosphorylates Ndc80, which promotes its binding to MTs.

**Fig 3.**
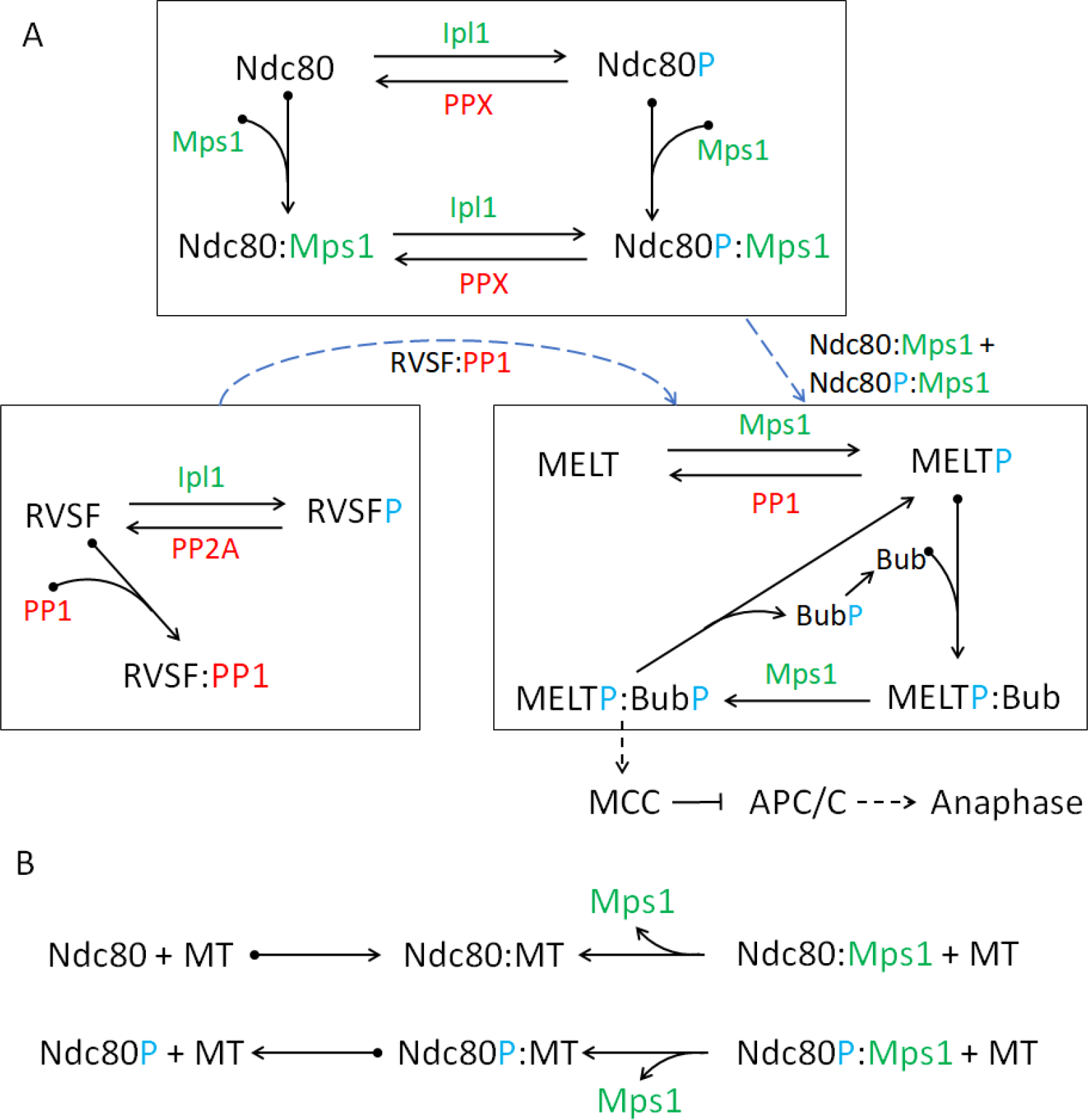
Molecular interactions at the KT. **(A)** Model of kinase (green) and phosphatase (red) activities at the KT, derived from the scheme in Fig. 2. The full scheme can be divided into three modules (shown within boxes), namely, Ndc80, MELT, and RVSF modules. See text for a detailed description of each module. ‘P’ (blue color) is used to depict the phosphorylation status of different motifs. Dashed arrows between the boxes show that Mps1 bound to either Ndc80 or Ndc80P phosphorylates MELT repeats, and PP1 bound to RVSF dephosphorylates them. **(B)** Scheme showing binding of Ndc80 to MT. The reactions on the first line show that Ndc80 (unphosphorylated) binds strongly to a MT, stabilizing the attachment, and Ndc80:Mps1 can also bind to a MT, which displaces Mps1 in an irreversible reaction. (MTs and Mps1 compete for the same binding site on Ndc80, therefore, upon MT binding Mps1 is removed from Ndc80.) We assume that Ndc80P (and Ndc80P:Mps1) can also bind to MTs (second line of reactions), but with a much larger dissociation constant, i.e., phosphorylation of Ndc80 promotes detachment of the MT from a KT.

The RVSF module contains different phosphorylation states and PP1-binding states of the RVSF motif on Spc105. By phosphorylating RVSF, Ipl1 prevents PP1 binding to this motif. PP2A opposes Ipl1 by dephosphorylating RVSF. There are five Spc105 molecules per KT (32), and one RVSF motif on each Spc105.

A third module contains different phosphorylation states and Bub-binding states of the six MELT motifs on Spc105 (32); i.e., 30 MELT motifs on each KT. This module is crucial for generation of the SAC signal. Mps1 phosphorylates MELT motifs and PP1 dephosphorylates them. In reality, phosphorylated MELT repeats bind Bub3-Bub1, and then phosphorylation of Bub1 by Mps1 allows binding of Mad1-Mad2 (33). The Mad1-Mad2 complex acts as template for generation of the Mitotic Checkpoint Complex (MCC), a diffusible signal that delays onset of anaphase by inhibiting the Anaphase Promoting Complex/Cyclosome (APC/C; a ubiquitin ligase) (34). We simplify this signaling cascade by lumping the SAC proteins into a single species called Bub, and assume that the state MELTP:BubP is capable of generating the MCC. The SAC signal is turned off by the dissociation of BubP from MELTP, and after dissociation BubP is dephosphorylated to Bub by some unknown phosphatase.

Coupling between the modules is shown with dashed arrows. Mps1 bound to either Ndc80 or Ndc80P phosphorylates the MELT repeats on Spc105, and PP1 bound to RVSF dephosphorylates MELT repeats. We assume that all substrates at the KT are accessible to their corresponding enzymes. For example, an Mps1 molecule bound to Ndc80 can phosphorylate all the available MELT repeats (30 of them), and the same holds true for Ipl1, PP2 and PP1 and their substrates.

Our model for attachment of Ndc80 to a MT is shown in Fig. 3B. KT-MT attachment is a complex process involving both the Ndc80 complex and the Dam1 complex (35). We focus only on the Ndc80 complex and assume that both phosphorylated and unphosphorylated forms of Ndc80 bind to a MT, and that the dissociation rate of Ndc80P:MT is much larger than that of Ndc80:MT, consistent with the observation that the affinity of Ndc80 for MTs decreases with the number of phosphorylations (9). It is also known that MTs and Mps1 compete for the same binding site on Ndc80 (25, 26). In our model, Mps1 is removed from Ndc80 upon MT binding.

### Kinetochore-Microtubule attachment dynamics

As mentioned earlier, in budding yeast, a MT attaches to a KT via Ndc80. To model the attachment dynamics we describe the attachment state of sister KTs by a two-dimensional vector (*m*, *n*), where the integers *m* and *n* correspond to the number of Ndc80s bound to a MT at each KT of a sister chromatid pair. Figure 4A shows the attachment dynamics of sister KTs. The symbols ‘s’ (red) and ‘a’ (green) correspond to syntelic and amphitelic attachments, respectively. A double arrow between states reflects that transitions can occur in both forward and backward directions. Figure 4B shows different realizations of the KT-MT attachment status as a function of time, calculated using the scheme in Fig. 4A. All the traces start in the syntelic attachment state and end in biorientation but, as shown later, in certain cases the KTs fail to reach biorientation.

**Fig 4.**
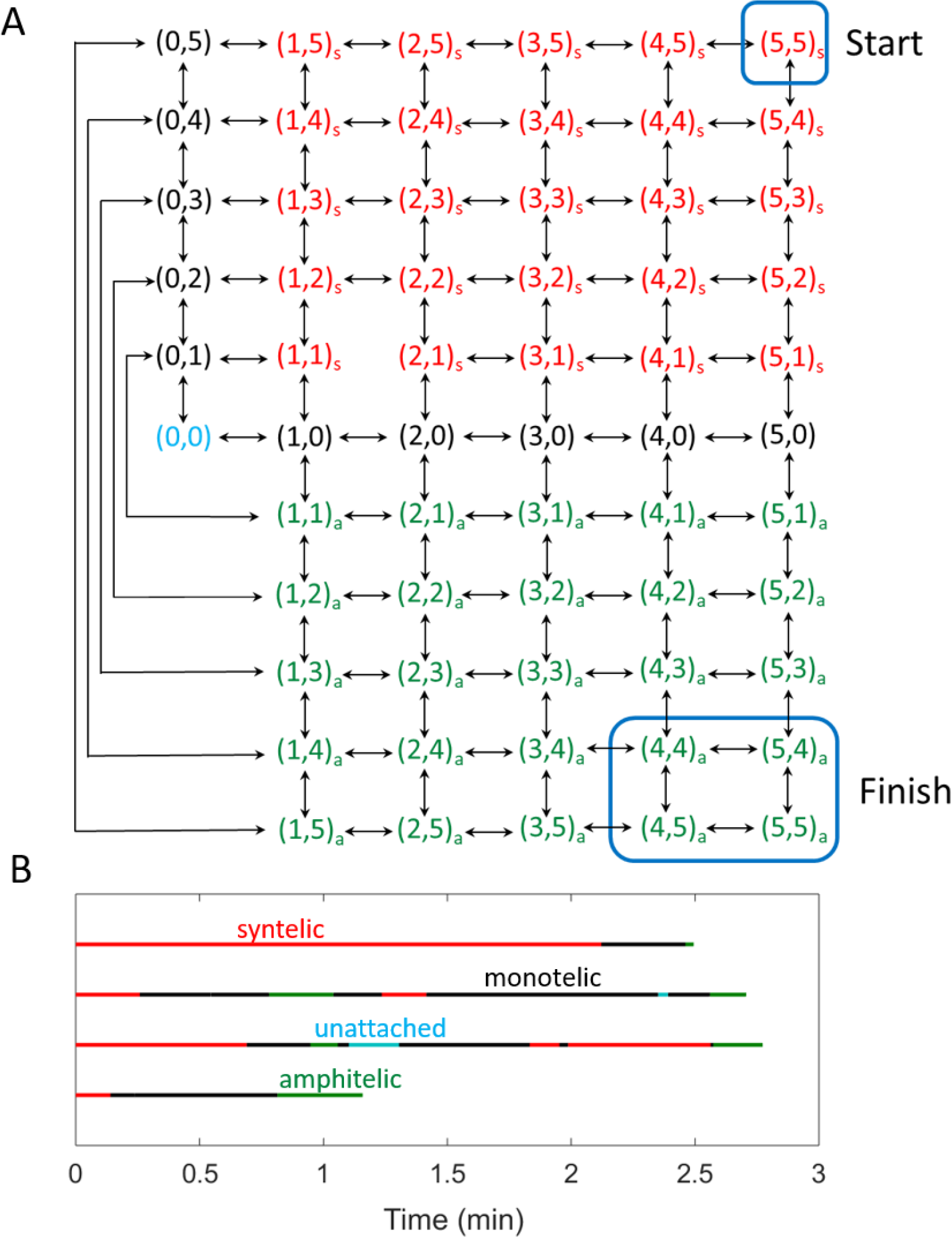
(A) A model of MT attachment to sister KTs through Ndc80. The vector (*m*, *n*) represents the number of Ndc80s bound to a MT at each KT. The attachment can be monotelic (black), syntelic (red with subscript s) or amphitelic (green with subscript a). The start and finish points of chromosome-alignment process are shown with blue boxes. We assume that the process stops (KTs reach biorientation) when the KTs spend more than 1 sec in the finish box without coming out. **(B)** Traces showing time-development of KT-MT attachment status calculated using the scheme in (A).

Since we are interested in understanding the dynamics of error correction, we choose that KTs start in a syntelic attachment state (5,5)_s_. This is reasonable as syntelic attachments are frequently observed during early mitosis in budding yeast (1, 2). As mentioned earlier, sister KTs attached to MTs from opposite spindle poles (i.e., amphitelic attachments) come under tension when the MTs exert forces (due to MT depolymerization) simultaneously on both KTs of a centromere. The typical time for such an event to occur has been estimated to be ~1 second (27). Thus, we assume that the amphitelic attachments come under tension when four or more Ndc80s (i.e., Ndc80+Ndc80P ≥ 4) are bound to the MTs on each side of a centromere for more than one second. In other words, the irreversible amphitelic-to-biorientation transition occurs when the system spends more than one second in the states (4,4)_a_, (5,4)_a_, (4,5)_a_, (5,5)_a_, without coming out.

For KTs in monotelic states (black color in Fig. 4), attachment of a MT to the unattached KT can be either amphitelic or syntelic (see Fig. 1). This attachment is a stochastic process that depends on factors like the rotational diffusion of sister chromatids in the monotelic state. We don’t account for the detailed motion of the chromosomes. Instead, we take a coarse-grained approach, introducing a parameter, P_syn_, to specify the ratio of transitions between monotelic → syntelic and monotelic → amphitelic states. For example, P_syn_ = 0.5 corresponds to the case where the KTs in monotelic state are equally likely to form syntelic or amphitelic attachments. Lower values of P_syn_ (< 0.5) corresponds to the case where KTs in a monotelic state are biased towards forming amphitelic attachments.

## Simulation

We prepared the model in an Excel file which contains the list of species, initial conditions, reactions corresponding to Ndc80, RVSF, MELT modules and KT-MT attachment, reaction-propensities, parameter values, and constraints. We wrote a MATLAB code which takes the Excel file as input and outputs another MATLAB file containing the stochiometric matrix and the propensity vector, which were used to prepare the code for Gillespie Stochastic Simulation of the model. The Excel and MATLAB files are provided in the online Supporting Information (SI). We stopped the simulation when one of the following two criteria was satisfied: (1) KTs reached biorientation, (2) time in simulation reached 10 mins. In budding yeast, the time interval between prometaphase to anaphase is approximately 15 mins (36). During that time the KTs must get bioriented and the SAC signal in the cytoplasm must be turned off (takes ~ 5 min). Thus 10 mins for reaching biorientation is a reasonable choice.

From our simulations we calculated the probability of biorientation within 10 min, fraction of time spent by KTs in different attachment states, average number of transitions between different states, and quantities related to SAC signal. The method used to calculate these quantities is described in SI. We performed 10000 stochastic simulations to calculate the statistics for different sets of parameter values. To study the case where formation of syntelic attachment is less probable than formation of amphitelic attachment the value of P_syn_ was chosen to be 0.1. Reducing P_syn_ to values smaller than 0.1 did not change the results significantly. P_syn_ = 0.5 was chosen to study the case where formation of syntelic and amphitelic attachments are equally probable. The activities of kinase and phosphatase are defined as (number of molecules)×(corresponding rate constant). For example, Ipl1 activity = Ipl1×kipl1. In our analysis we keep the number of molecules of Ipl1, PPX, PP2A constant (all equal to one), and change their activities by changing their corresponding rate constant.

## Results

### Kinase-phosphatase balance during error correction

Here we explore how Ipl1 and PPX activities affect the probability of biorientation. For this analysis we use the Ndc80-MT attachment model shown in Fig. 4 and choose P_syn_ = 0.1. First, we determine what happens in the absence of kinase and phosphatase activities. To this end, we set phosphorylation/dephosphorylation rate of Ndc80 by Ipl1/PPX to zero (kipl1 = kppx = 0) and calculate the dependence of biorientation probability on kdndc, the Ndc80:MT dissociation rate. In this case kdndc1 (Ndc80P:MT dissociation rate) becomes unimportant because the Ndc80s start in the unphosphorylated state and are never phosphorylated. We find that biorientation probability attains a peak value of one at kdndc = 0.7 sec^−1^ (Fig. 5). Thus, if the dissociation rate is properly tuned then error correction and biorientation (within 10 mins) can, in principle, occur even in the absence of kinase and phosphatase activities.

**Fig 5.**
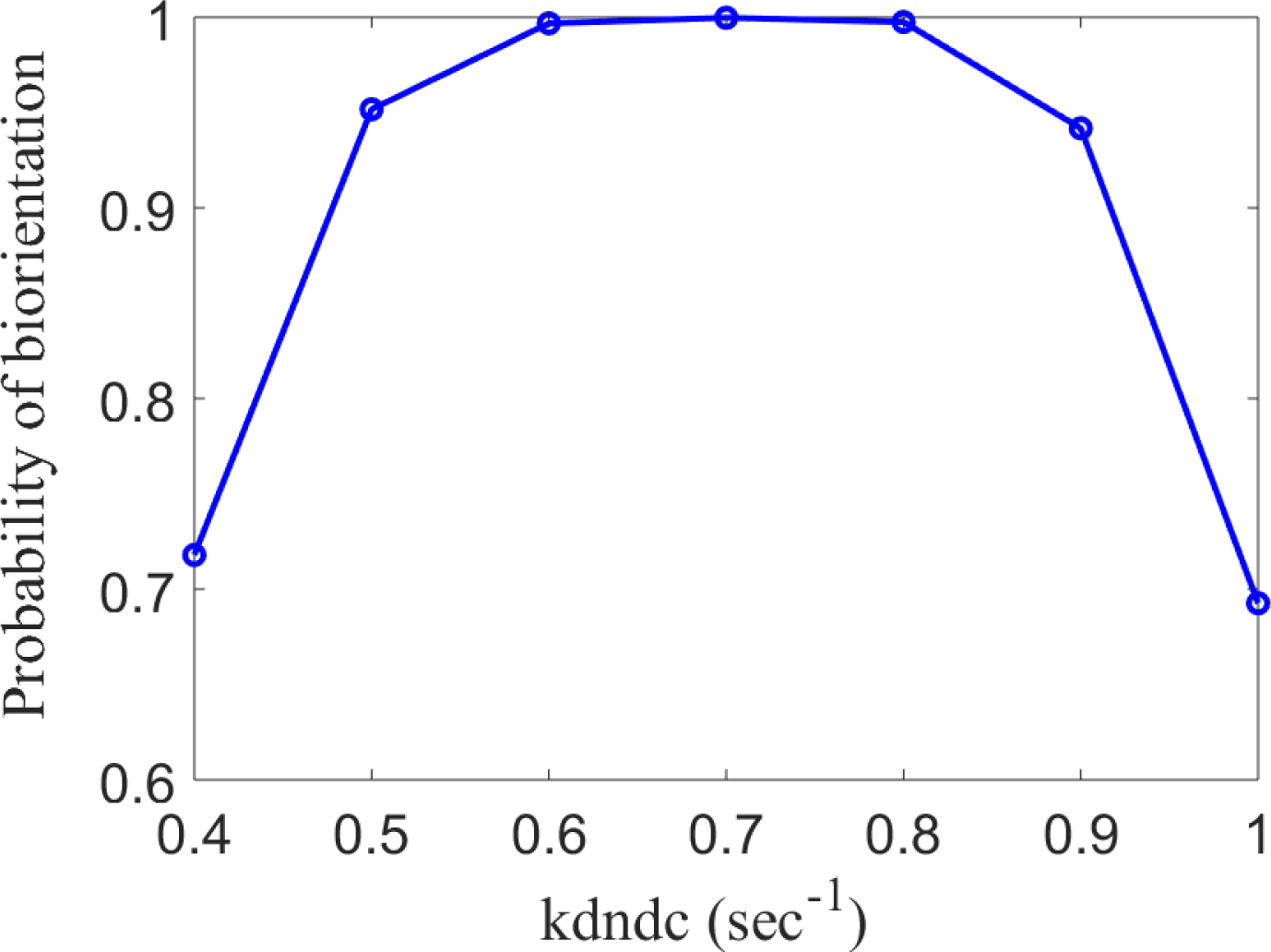
Probability of biorientation in absence of kinase and phosphatase activities. Probability of sister KTs reaching biorientation within 10 mins, starting from the syntelic attachment state. For this analysis we chose kipl1 = 0 and kppx = 0. The probability of biorientation for kdndc = 0.7 sec^−1^ is one. Thus, error correction and biorientation can in principle occur even in the absence of kinase and phosphatase activities.

Since Ipl1 activity is required for error correction (3–5), the value of kdndc is probably less than 0.7 sec^−1^. Indeed, in-vitro measurements of dissociation rate show that in the absence of Dam1 complex, kdndc = 0.44 sec^−1^ and in the presence of Dam1 complex it drops down to 0.23 sec^−1^ (37). If the value of kdndc1 is also less than 0.7 sec^−1^, biorientation cannot occur efficiently (see SI). This suggests that kdndc and kdndc1 are smaller and greater than 0.7 sec^−1^, respectively.

The probability of biorientation when kdndc is smaller and kdndc1 is larger than 0.7 sec^−1^ (kdndc = 0.1 sec^−1^ and kdndc1 = 1.5 sec^−1^) is shown in Fig. 6. Different curves correspond to different values of kppx. For each curve the biorientation probability peaks over a range of kipl1 values. As the phosphatase activity is increased (by increasing kppx), the peak occurs at higher values of kipl1, which shows that a ‘balance’ between the two activities is required. The peak value of probability in each case is one, implying that the chance of not reaching biorientation is less than 1/10000. Note, the range of Ipl1 activity over which the biorientation probability is high (close to one) increases with kppx.

**Fig 6.**
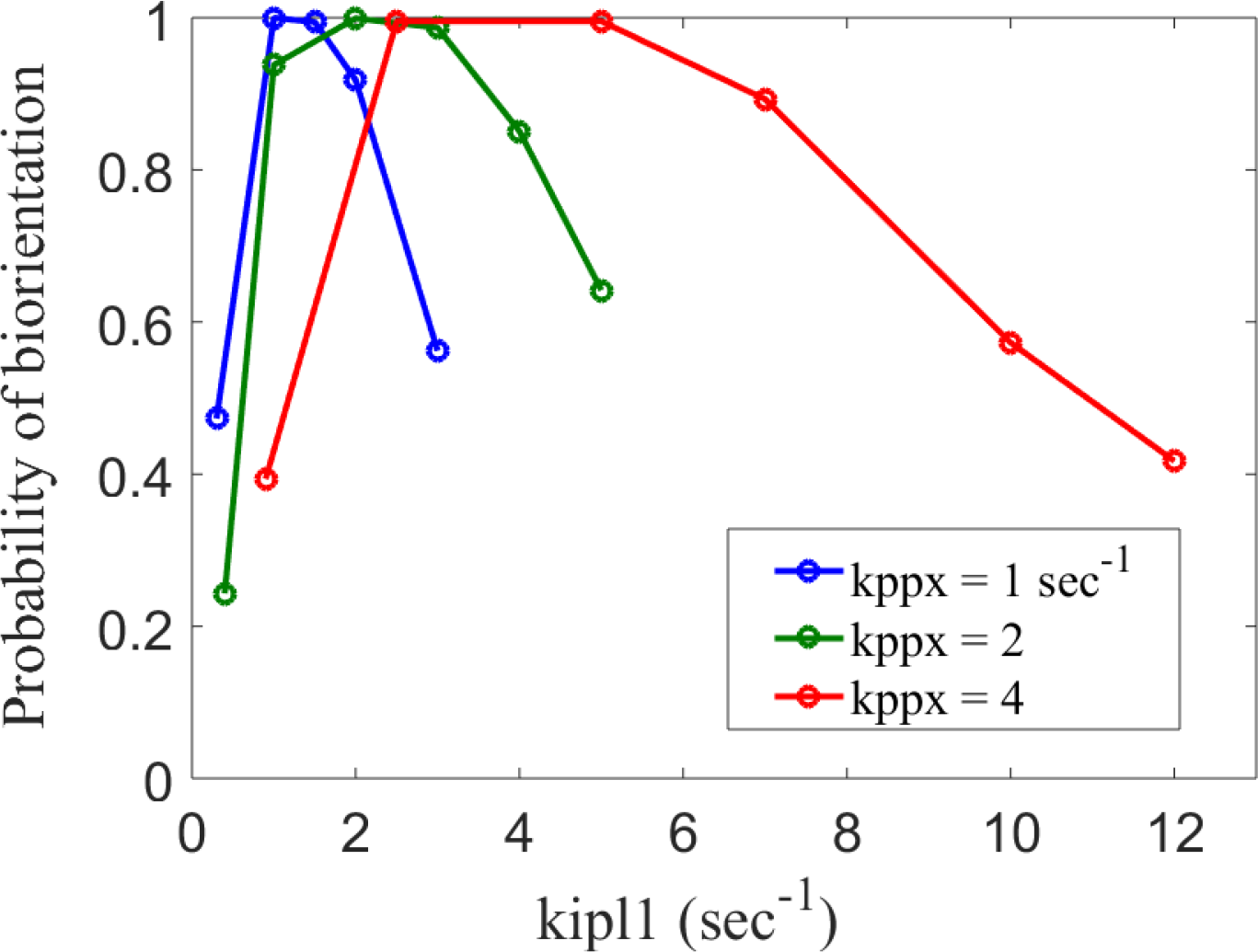
Kinase phosphatase balance. Probability of sister KTs reaching biorientation within 10 mins, starting from the syntelic attachment state. For this analysis we chose kdndc = 0.1 sec^−1^ and kdndc1 = 1.5 sec^−1^. Different curves correspond to different kppx (phosphatase activity) values. Peak in biorientation probability shifts to higher values kipl1 with increasing kppx, which suggests that a balance between the kinase and phosphatase activities is required for biorientation to occur efficiently.

To quantify the dynamics of KT-MT attachment, we calculated the fraction of time spent by sister KTs in different attachment states and the average number of transitions between different attachment states in a simulation run. The method to calculate these quantities is described in Section 5 of SI. The statistics of attachment dynamics for the case kppx = 4 sec^−1^ (red curve in Fig. 6) is shown in Table 1. The balance point between kinase and phosphatase activities occurs at kipl1 = 4 sec^−1^. Below the balance point (kipl1 = 1 sec^−1^), the biorientation probability drops because the KTs fail to correct the initial syntelic attachment quickly. This can be seen in the large fraction of time spent by KTs in the syntelic attachment state (92%). Above the balance point (kipl1 = 7 sec^−1^), the biorientation probability drops because the Ndc80-microtubule attachments, even the correct ones are disrupted too often. This can be seen in the increased fraction of time spent in the unattached at monotelic states as well as the increased number of amp → mon and mon → unattached transitions (Table 2).

**Table 1.** Fraction of time spent by KTs in different KT-microtubule attachment states as a function of Ipl1 activity.

|  |  |  | Percent of time in each state |  |  |  |
| --- | --- | --- | --- | --- | --- | --- |
| $k_{ip11} (\text{sec}^{-1})$ | $k_{ppx} (\text{sec}^{-1})$ | Pbior | syntelic | unattached | monotelic | amphitelic |
| 1 | 4 | 0.5 | 92 | 1 | 6 | 1 |
| 4 | 4 | 1 | 39 | 5 | 34 | 22 |
| 7 | 4 | 0.9 | 15 | 14 | 45 | 26 |

**Table 2.** Mean number of each type of transition as functions of Ipl1 activity. The states are monotelic (mon), amphitelic (amp), syntelic (syn), unattached (unatt.).

| $k_{ip11} (\text{sec}^{-1})$ | $k_{ppx} (\text{sec}^{-1})$ | $\text{mon} \rightarrow \text{syn}$ | $\text{amp} \rightarrow \text{mon}$ | $\text{mon} \rightarrow \text{unatt.}$ |
| --- | --- | --- | --- | --- |
| 1 | 4 | 0.03 | 0.02 | 0.02 |
| 4 | 4 | 0.19 | 0.75 | 0.54 |
| 7 | 4 | 0.63 | 4.7 | 4.2 |

We also determined the dependence of attachment dynamics on the absolute value of activities. Table 3 shows the fraction of time in different attachment states at different kinase-phosphatase values (near the optimal value of Ipl1 activity). As the values of kinase and phosphatase activities are increased, the time spent in the initial syntelic attachment drops and the time spent in unattached, monotelic and amphitelic states increases marginally. The average time for biorientation was 1.9 mins. The distribution of biorientation times is shown in Fig. S2 (see SI).

**Table 3.** Statistics of the KT-MT attachment transitions as a function of Ipl1 activity. The probability of biorientation in each case is one. As the balance point of activities is increased, the time spent in syntelic attachment state drops.

| kip11 ($\text{sec}^{-1}$) | kppx ($\text{sec}^{-1}$) | $\langle \text{Time} \rangle$ (min) | Percent of Time in each state | | | |
| --- | --- | --- | --- | --- | --- | --- |
|  |  |  | syntelic | unattached | monotelic | amphitelic |
| 1 | 1 | 1.9 | 50 | 3 | 30 | 17 |
| 2 | 2 | 1.9 | 44 | 4 | 32 | 20 |
| 4 | 4 | 1.9 | 39 | 5 | 34 | 22 |

In summary, error correction and then biorientation occurs efficiently when kinase and phosphatase activities are balanced. A balance at higher absolute values of kinase and phosphatase activities has two advantages. First, it increases the range over which error correction and then biorientation can occur with high probability. Second, it increases the fraction of time spent in the monotelic and unattached states. As shown later, this marginally improves the SAC signaling strength.

### Analytical calculation of balance point of kinase and phosphatase activities

In the previous section we found that a balance between kinase and phosphatase activities is needed, but it was not clear what the balance means quantitatively. Here we determine the condition that defines the balance between the two activities. To do so, the following reasoning is used: in the absence of kinase and phosphatase activities, biorientation occurs efficiently when kdndc = 0.7 sec^−1^. Then in the general, when kinase and phosphatase activities are nonzero, the activities should balance to produce an effective dissociation rate of 0.7 sec^−1^. Thus, the balance condition can be determined by calculating the effective dissociation rate of Ndc80-MT attachment and setting it to 0.7 sec^−1^.

To calculate the effective dissociation rate of Ndc80-MT attachment (k_eff_), we use the scheme shown below.

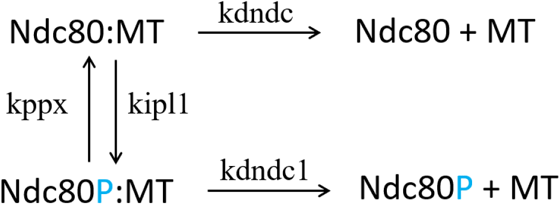

Dissociation of Ndc80 from MT can occur from the states Ndc80:MT or Ndc80P:MT. If we define the rate of dissociation as the inverse of mean first passage time, then the dissociation rate starting from the state Ndc80:MT is given by (see SI)

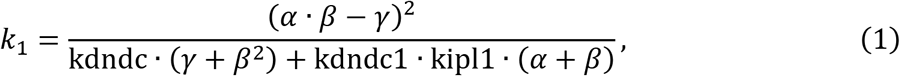

where *α* = kdndc + kipl1, *β* = kdndc1 + kppx, and γ = kipl1 ∙ kppx. In the above expression we omit the number of molecules Ipl1 = PPX = 1. Similarly, the rate of dissociation starting from Ndc80P:MT state is given by

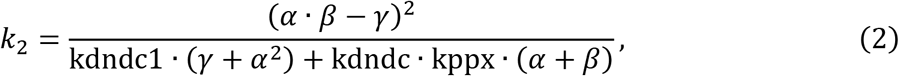

Assuming phosphorylated and dephosphorylated states of Ndc80 are in equilibrium, the effective dissociation rate of Ndc80-MT attachment can be written as

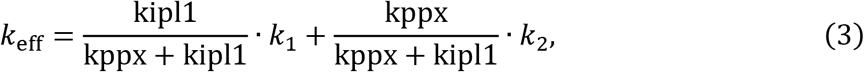

If we assume that phosphorylation/dephosphorylation rates are much larger than the two dissociation rates, i.e., *α* ≈ kipl1 and *β* ≈ kppx, then the effective dissociation rate simplifies to

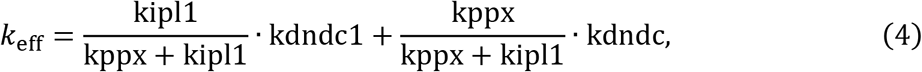

and the ratio between kinase and phosphatase activities can be written as

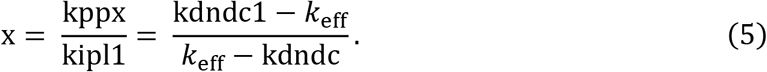

Substituting the values kdndc = 0.1 sec^−1^ and kdndc1 = 1.5 sec^−1^, and k_eff_ = 0.7 sec^−1^ in the above equation, we find x = 1.33 which is close to our observation in Fig 5, that Ipl1 and PPX activities must be approximately equal for error correction to occur efficiently. Fig 7 shows the curves at which the effective dissociation rate is equal to 0.6, 0.7, 0.8 sec^−1^. For these values the biorientation probability shown in Fig 6 is high. The dashed double arrow shows the range of kipl1 over which biorientation can occur efficiently. As observed earlier, this range increases with kppx.

**Fig 7.**
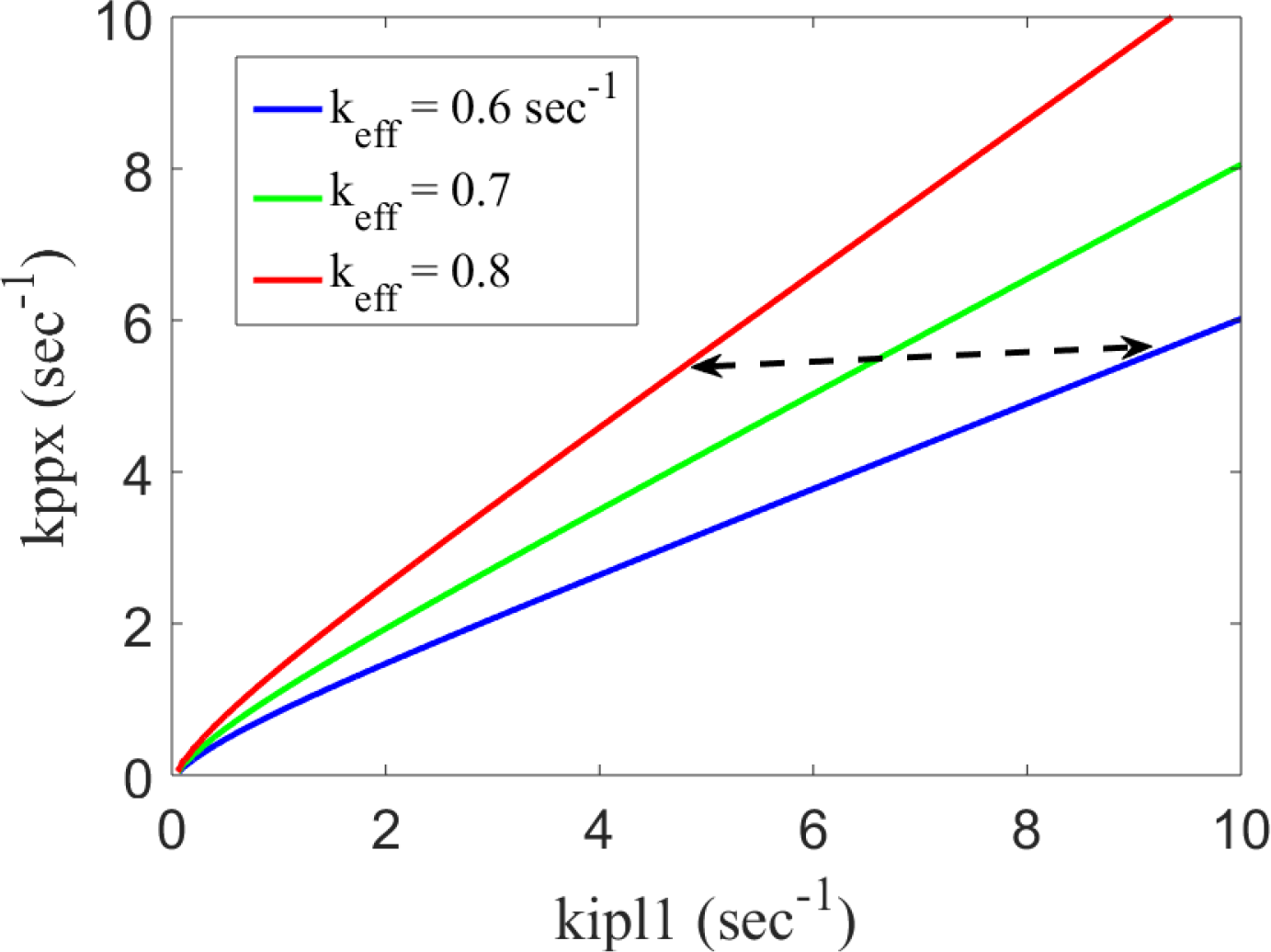
Constant value curves for effective MT-Ndc80 dissociation rate. Curves on which the effective dissociation constant k_eff_ is constant. The curves were calculated using Eq. 3. At a fixed value of kppx the interval between the red and the blue curve is the range of kipl1 over which the biorientation probability is high (see Fig. 5). This range increases with increasing kppx.

### Effect of kinetochore orientation on biorientation of KTs

Along with the Ipl1-dependent method of error correction, geometrical factors can also contribute towards biorientation of KTs. It has been proposed that MT attachment from one spindle pole orients the sister KT in a way that attachment from the opposite spindle pole becomes more probable (38). Such geometrical orientation of KTs can promote biorientation by reducing the chances of formation of syntelic attachments. We asked what are the relative contributions of Ipl1-dependent MT-KT detachment and KT geometrical orientation to achieving biorientation.

To study the effect of geometrical orientation we changed the parameter, P_syn_, which was used to specify the ratio of transition between monotelic → syntelic and monotelic → amphitelic states. In previous analysis we chose P_syn_ = 0.1; here we compare it with P_syn_ = 0.5, representing the case in which there is no geometrical bias preventing the formation of syntelic attachments, i.e., the transition probabilities from monotelic to syntelic and amphitelic states are equal.

The comparison between the probability of biorientation for P_syn_ = 0.5 and 0.1 is shown in Fig 8. For P_syn_ = 0.5, at Ipl1 activity = 4 sec^−1^, the biorientation probability is approximately 0.9614, which shows that if the kinase and phosphatase activities are balanced, then orientation of sister KT is not critical and Ipl1-dependent error correction of syntelic attachments is good enough to reach an accuracy of 0.9614 (~ 96%).

**Fig 8.**
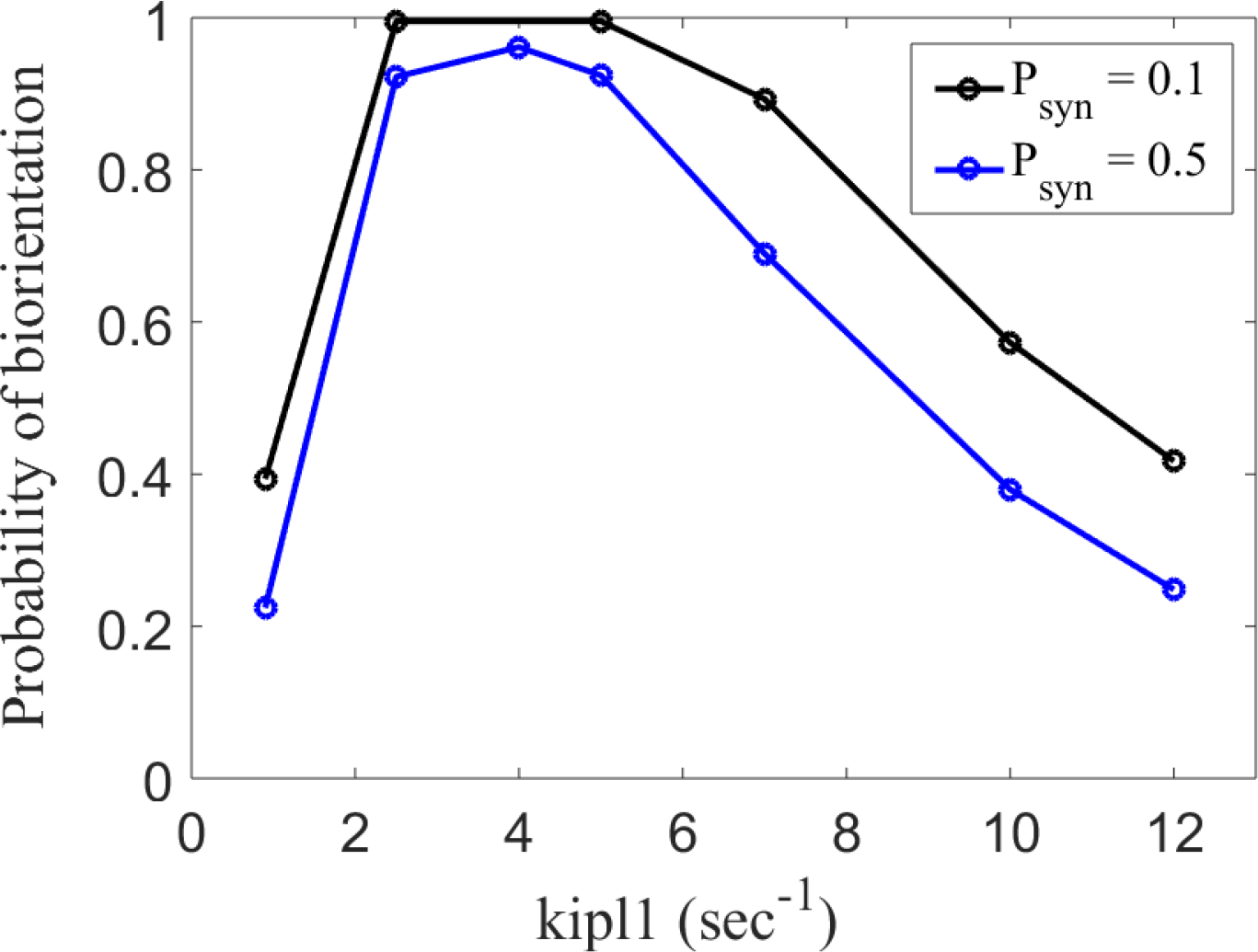
Effect of geometrical orientation of KTs on biorientation probability. Probability of biorientation within 10 mins for different values of P_syn_. KTs start in syntelic attachment state. P_syn_ = 0.5 corresponds to the case in which KTs in monotelic state transition to syntelic and amphitelic states with equal probability. P_syn_ = 0.1 corresponds to the case in which KTs in monotelic state transition to amphitelic state 10 times more often than syntelic state.

To understand why geometrical orientation of KTs is not that important we calculated the statistics of attachment dynamics for P_syn_ = 0.5 case (see Table 4). As expected, the average number of monotelic to syntelic transitions increases. The average biorientation time for the realizations in which the KTs reach biorientation was approximately 3.2 mins. The average number of times the KTs were in the syntelic attachment state was 2.7. This is because the starting syntelic state and mon → syn transitions contribute 1 and 1.7, respectively. Using these numbers and the quantities in Table 3, we find that the average of the time interval between entering and exiting syntelic attachment state is 3.2 min×(46/100)/2.7 = 0.54 min, which is much smaller than 10 mins. This shows that in absence of geometrical orientation of KTs the probability of biorientation does not drop significantly because when the kinase-phosphatase activities are balanced, the KTs have enough time to correct additional syntelic attachments.

**Table 4:** Statistics of attachment process for P_syn_ = 0.5. Top table shows the fraction of time spent in different attachment states. The bottom table shows the average number transition between different attachment states during single syntelic → biorientation process. The numbers in bold are the values for P_syn_ = 0.1 case. The abbreviations are monotelic (mon), amphitelic (amp), syntelic (syn).

|  |  |  | Percent of Time in each state |  |  |  |
| --- | --- | --- | --- | --- | --- | --- |
| kipl1 (sec<sup>-1</sup>) | kppx (sec<sup>-1</sup>) | Pbior | syntelic | unattached | monotelic | amphitelic |
| 4 | 4 | 0.9610 (**1**) | 46% (**39**) | 5% (**5**) | 34% (**34**) | 15% (**22**) |

**Table 4:** Statistics of attachment process for $P_{\text{syn}} = 0.5$ .
| kipl1 (sec<sup>-1</sup>) | kppx (sec<sup>-1</sup>) | mon → syn | amp → mon | mon → unattached |
| --- | --- | --- | --- | --- |
| 4 | 4 | 1.7 (**0.19**) | 0.73 (**0.75**) | 1.12 (**0.54**) |

### Coupling between error correction and SAC

Here we analyze the coupling between error correction and SAC. The coupling is quantified by the strength of SAC signal, and PP1 and Mps1 binding. These quantities are defined as the average (over time) occupancy of the corresponding states, i.e, NSAC = MELTP:BubP, NPP1 = RVSF:PP1, and NMps1 = Ndc80:Mps1+Ndc80P:Mps1. The method for calculating these quantities is described in SI. First, to quantify a strong SAC signal and to determine the value of parameter kpp1 (the dephosphorylation rate of MELTP by PP1) we compare our simulation results with experimental data.

Bubs bind to MELT motifs to initiate SAC signaling. The number of Bubs needed to generate a strong SAC signal is not known. In an experiment where the PP1 activity was suppressed, it was found that approximately 20 (~10 on each KT) Bubs bound to a single pair of unattached KTs (32), so we use this number as the reference point for a strong SAC signal. We calculated the strength of SAC signal in the absence of PP1 activity by setting the forward rate of PP1 binding to zero (kfpp1 = 0). In this case Bub binding was NSAC = 18.6, which is close to the reference value. In our simulation this number is determined by the concentration of Bub, the association/dissociation rate of Bub to MELT, and the phosphorylation rate of MELT and Bub by Mps1 (see SI for these parameter values).

Next, we simulated the mutant Spc105-RVAF (21), to determine the value of kpp1. In this mutant Ipl1 cannot phosphorylate RVSF motif and therefore PP1 binds to the RVSF motif constitutively. Despite unhindered binding of PP1, the SAC was not turned off prematurely (21). This mutant was simulated by setting the phosphorylation rate of RVSF motif to zero (kipl1a = 0). Table 4 below shows NSAC, NPP1, and NMps1 for different values of kpp1. For kpp1 = 1/100 sec^−1^, the SAC signal strength is close to the reference value. Thus, we choose kpp1 = 1/100 sec^−1^. The strength of the SAC signal is determined by the competition between Mps1 and PP1 activities. Mps1 activity (kmps1×NMps1) is 1.1 sec^−1^ and PP1 activity is 0.036 sec^−1^. That is, the Mps1 activity is approximately 30 times larger than PP1 activity.

Having defined the reference value for a strong SAC signal and determined the value of kpp1, next, we quantify the coupling between the SAC and error correction. Table 5 shows NSAC, NPP1 and NMps1 for different values of kipl1 (corresponding to the red curve in Fig. 6). NPP1 is constant because it does not depend on kipl1 and kppx; its value is set by kipl1a and kpp2, the phosphorylation/dephosphorylation rates of RVSF motif by Ipl1 and PP2A, respectively. At kipl1 = 1 sec^−1^, NSAC is considerably smaller than the reference value (of 20) because the KTs get stuck in the syntelic attachment state with most of the Ndc80s bound to MT. This precludes the binding of Mps1 and activation of SAC. As kipl1 is increased to 4 sec^−1^, the fraction of time spent by the sister KTs in the unattached and monotelic states increases (see Table 1). This leads to higher binding of Mps1 (NMps1) and a significantly stronger SAC signal. At kipl1 = 7 sec^−1^, NMps1 and NSAC increase further but, as shown earlier, the probability of biorientation of KTs drops.

**Table 5:**
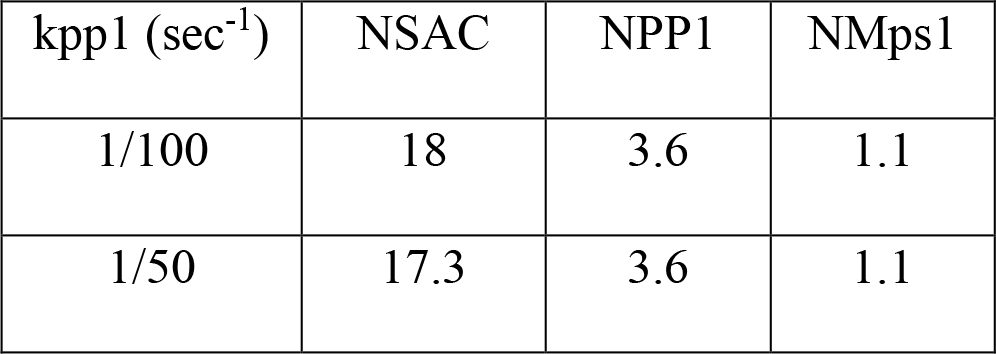
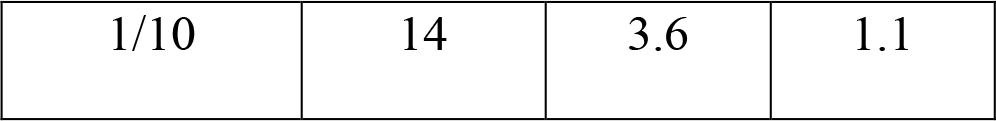
Average number of SAC signaling state, and PP1 and Mps1 bound to KT during as a function of kpp1, the PP1 dependent dephosphorylation rate of MELT motifs. The data presented in the Table was used to set the value of kpp1 to 1/100 sec^−1^.

| $k_{pp1} (\text{sec}^{-1})$ | NSAC | NPP1 | NMps1 |
| --- | --- | --- | --- |
| 1/100 | 18 | 3.6 | 1.1 |
| 1/50 | 17.3 | 3.6 | 1.1 |
| 1/10 | 14 | 3.6 | 1.1 |

We also determined how the strength of SAC signal depends on the absolute values of kinase-phosphatase activities. The values of NSAC, NPP1 and NMps1 at different value of activities (while maintaining the balance) are given Table 6. Earlier, we found that a balance point at higher activity leads to KTs spending a higher fraction of time in the unattached and monotelic states. Here we see that it leads to a small increase in NMps1 and NSAC.

**Table 6:** Average number of SAC signaling state, PP1 molecules bound to KT, and Mps1 molecules bound to Ndc80 as function of Ipl1 dependent phosphorylation rate of Ndc80 (kipl1).

| kipl1 (sec<sup>-1</sup>) | kppx (sec<sup>-1</sup>) | NSAC | NPP1 | NMps1 |
| --- | --- | --- | --- | --- |
| 1 | 4 | 8.4 | 3.7 | 0.1 |
| 4 | 4 | 18.5 | 3.6 | 1.2 |
| 7 | 4 | 23.1 | 3.6 | 1.8 |

**Table 7:** Average number of SAC signaling state, PP1 molecules bound to KT, and Mps1 molecules bound to Ndc80 at different values of kinase and phosphatase activities.

| kip11 ($\text{sec}^{-1}$) | kppx ($\text{sec}^{-1}$) | NSAC | NPP1 | NMps1 |
| --- | --- | --- | --- | --- |
| 1 | 1 | 16.3 | 3.6 | 0.9 |
| 2 | 2 | 16.8 | 3.6 | 1 |
| 4 | 4 | 18.5 | 3.6 | 1.2 |

## Discussion

Fidelity of chromosome segregation process is guarded by two coupled mechanisms: error correction in KT-MT attachments and the SAC. The error correction mechanism removes erroneous attachments between KT and MT, and the SAC ensures that cells do not proceed to anaphase until all chromosomes are correctly attached. In this paper we present a stochastic model to study how the opposing activities of these kinases and phosphatases affect these two mechanisms in budding yeast. Our model includes the dynamics of MT attachment to KT through Ndc80, binding of key kinases PP1 and Mps1 to the KT and the activities of Ipl1 and PP2A and PPX (an unknown phosphatase that opposes Ipl1). We used this setup to calculate the probability that a pair of KTs reach biorientation within 10 mins, starting from the syntelic attachment state, and also the statistical details of the attachment process, like the time spent by KTs in different attachment states and the number of transitions between different attachment states.

We find that a balance between the kinase (Ipl1) and phosphatase (PPX) activities is required for KTs to reach biorientation efficiently. The balance point is defined by the ratio between the kinase and phosphatase activities. If the Ipl1 activity is below the balance point, biorientation probability drops because MT-Ndc80 attachments are stabilized excessively and correcting initial syntelic attachments takes longer time. On the other hand, if Ipl1 activity is above the balance point, the correct MT-Ndc80 attachments are destabilized too often for the KTs to reach biorientation efficiently. We derive an approximate analytical formula that defines the balance point and show that at higher absolute value of kinase and phosphatase activities (while maintaining the balance) the range of over which balance can be achieved is larger. This can make the error correction process more robust to fluctuations in absolute value of kinase and phosphatase activities.

The error correction process generates unattached kinetochores which then can initiate SAC signaling. This is one way the error correction mechanism is coupled to the SAC. Our analysis of this coupling shows that to maintain a strong SAC signal, first, the Ipl1 activity must be equal to (or larger than) the value defined by the balance point. Otherwise the SAC strength drops because the KTs get stuck in the syntelic attachment state. And second, the activity of Mps1 must be significantly larger than PP1 activity (our estimate is 30 time larger). Otherwise, the dephosphorylation of MELT motifs by PP1 starts reducing the signal strength. The strength of SAC signal crucially depends on the Mps1 activity. When the KTs get stuck in the syntelic attachment state, the SAC signal drops because KTs cannot recruit enough Mps1. Interestingly, experiments show that even after biorientation some residual Mps1 remains on the KTs (25). If this residual Mps1 is present on KTs in the syntelic attachment state, it can probably initiate SAC signaling. This pathway of initiating SAC would not depend on the creation of unattached kinetochores.

When one KT attaches to one spindle pole, the sister KT is constrained to face the opposite spindle pole. We calculated how the probability of biorientation is affected when that constraint is relaxed and found that if the Ipl1 activity is near its optimal value (as determined by our analysis) then the Ipl1-dependent error correction mechanism is sufficient for achieving timely biorientation with 96% accuracy, regardless of geometric constraints. However, such constraints can modestly improve the efficiency of reaching biorientation.

Several questions still remain unanswered. First, what is the role of PP2, if at all there is one? The phosphatase PP2A dephosphorylates RVSF motif to facilitate PP1 binding. However, experiments show that dynamic regulation of PP1 binding to Spc105 is not essential for mitosis (21). Therefore, the role of PP2A in budding yeast also seems unimportant. Second, how PP1 turns off the SAC signal? PP1 can actively promote dissociation of Bubs, i.e., the dissociation rate is proportional to PP1; or PP1 can oppose the binding of Bubs by dephosphorylating the MELT repeats (as in our model). In the latter case PP1 does not actively turn off the SAC, but only prevents its activation. Another interesting possibility is that PP1 dephosphorylates the Bub in MELTP:BubP to turn off the SAC. We think our model will provide a suitable starting point for analyzing these different possibilities when more experimental data becomes available. We also think our model can be adapted to study the same questions in fission yeast and mammals. In these organisms the number of binding sites for kinases and phosphatases at KT, and MT attachments per KT is much larger. Furthermore, in these organisms merotelic attachment states are observed. Including these details in the model will significantly increase the number of species and hence the computational cost, but nevertheless will provide a method to study error correction and its coupling to SAC in these organisms.

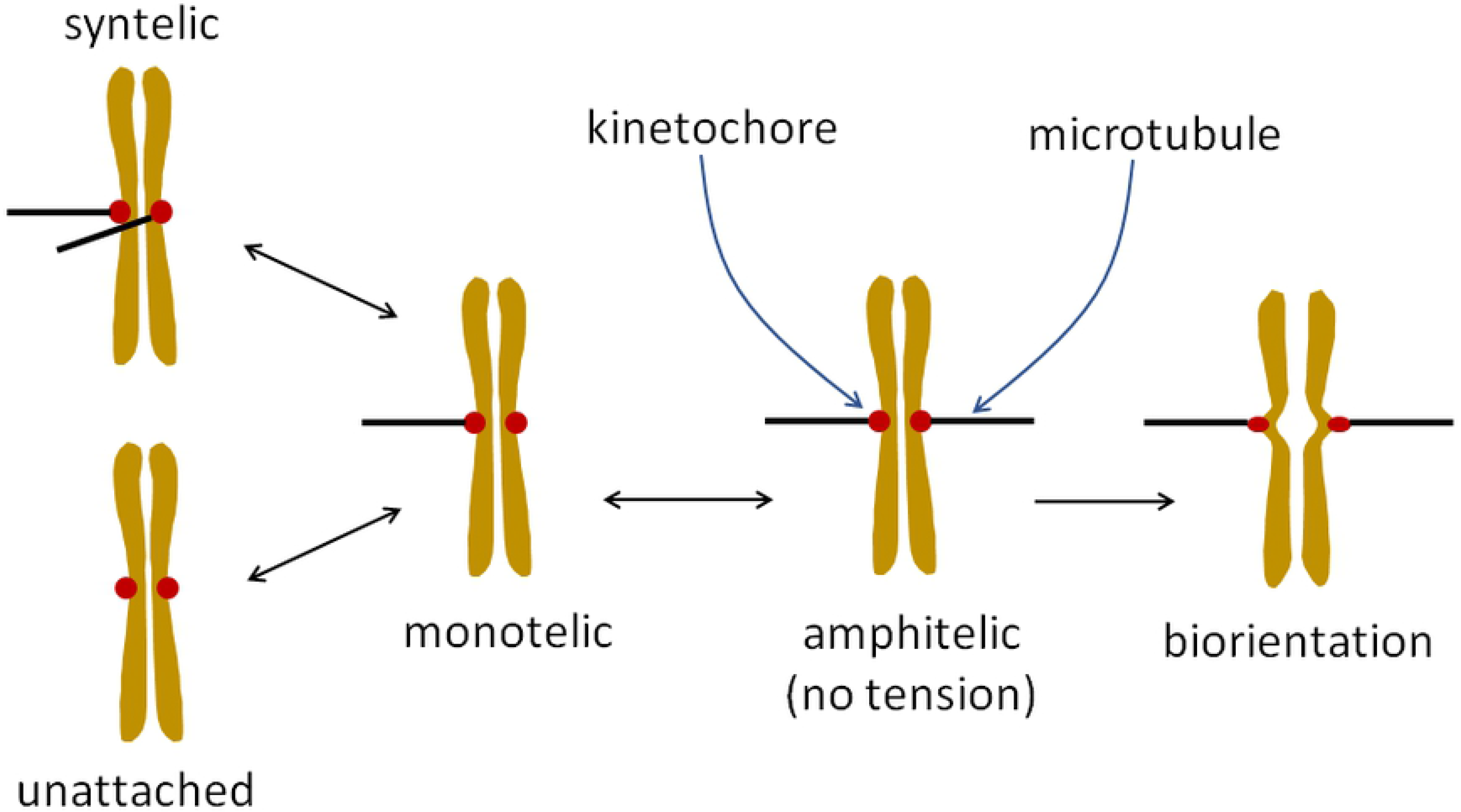

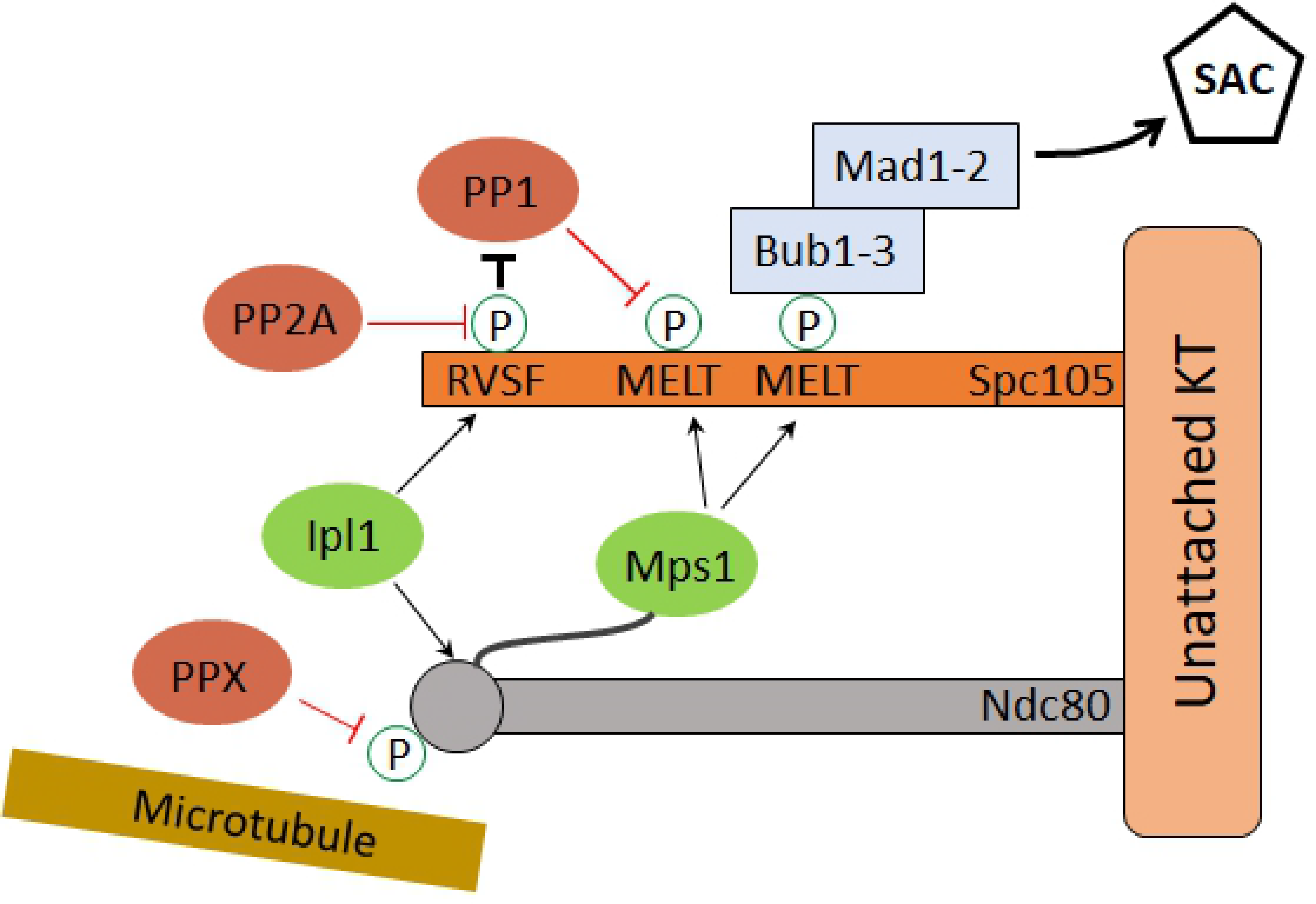

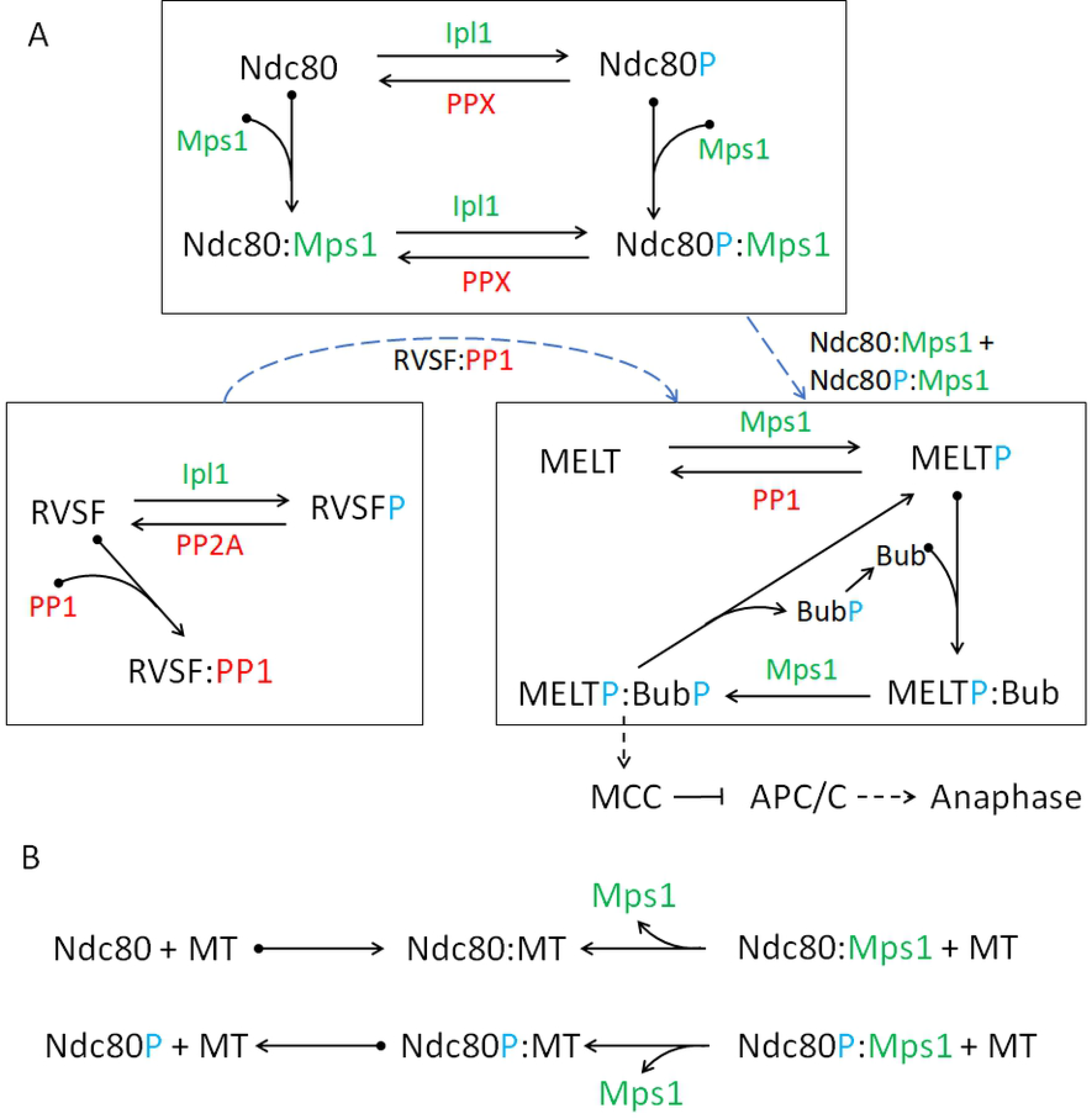

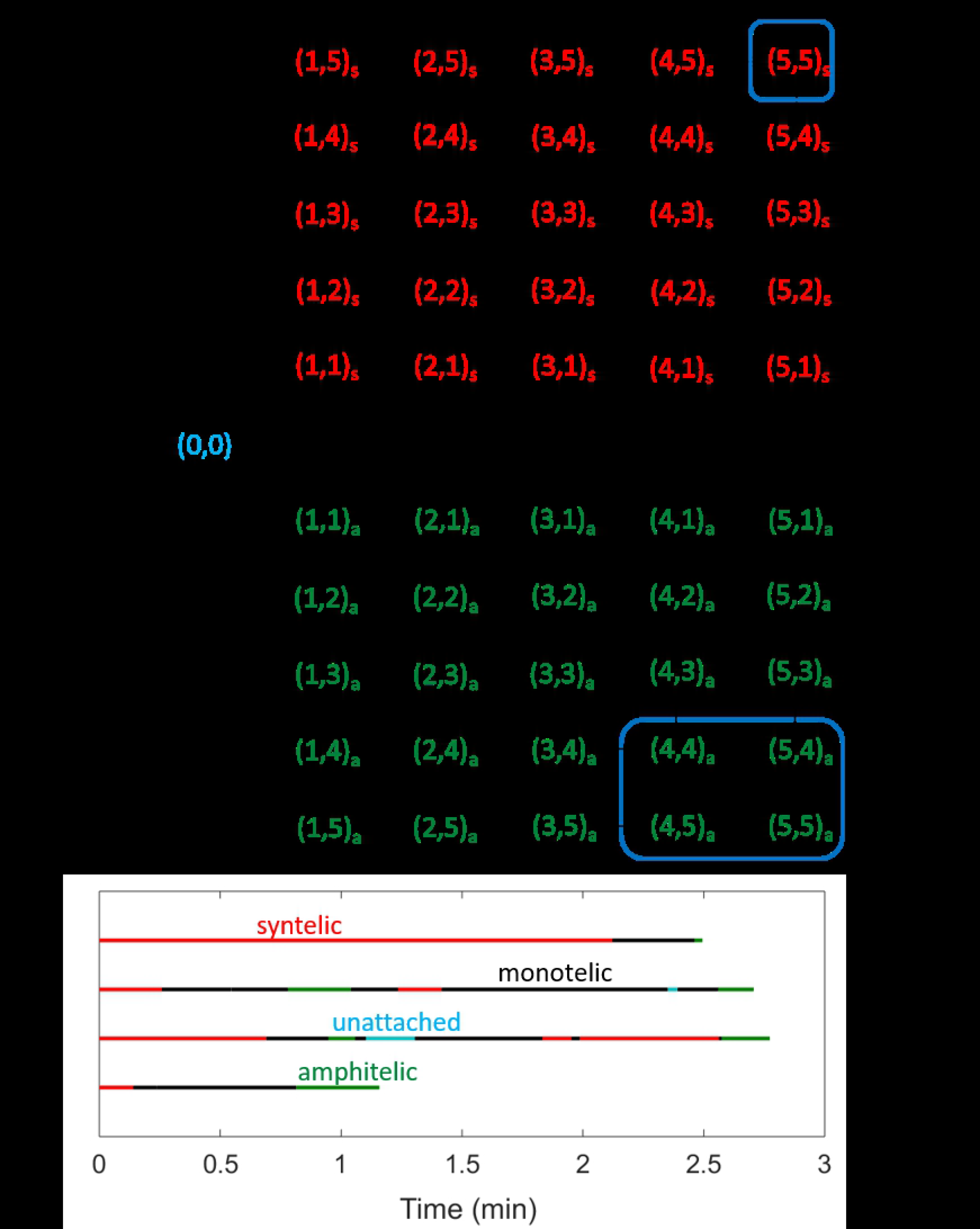

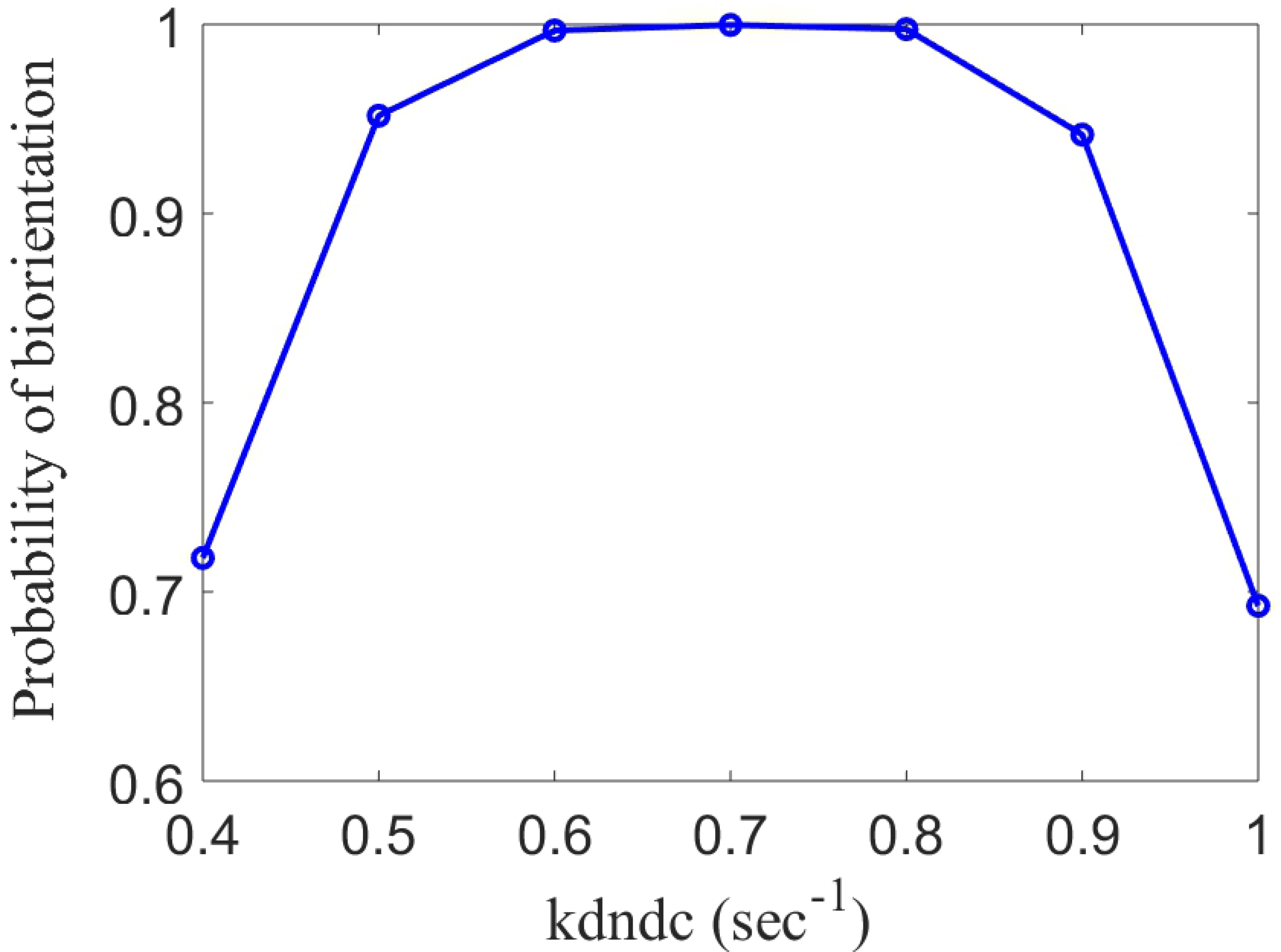

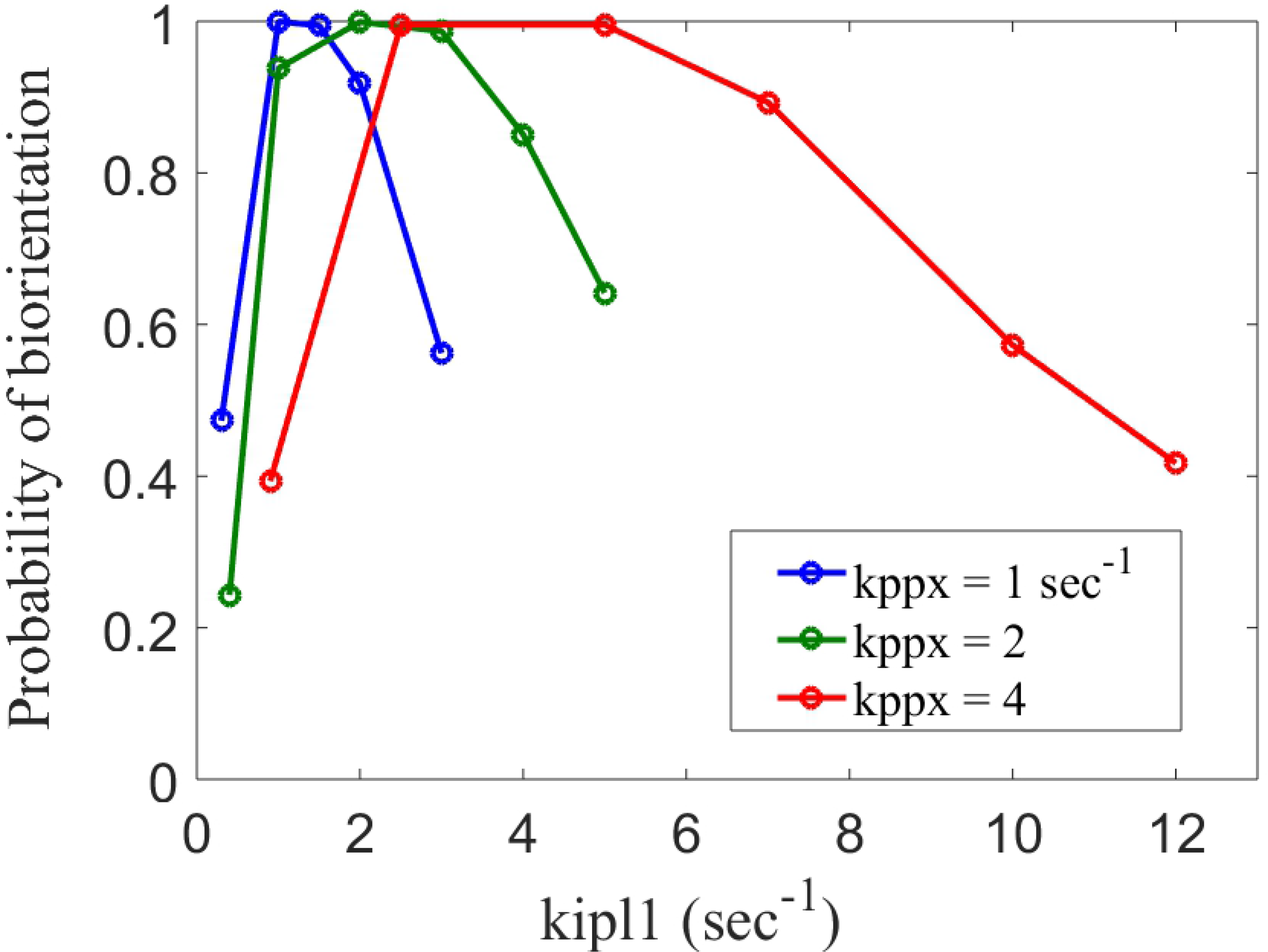

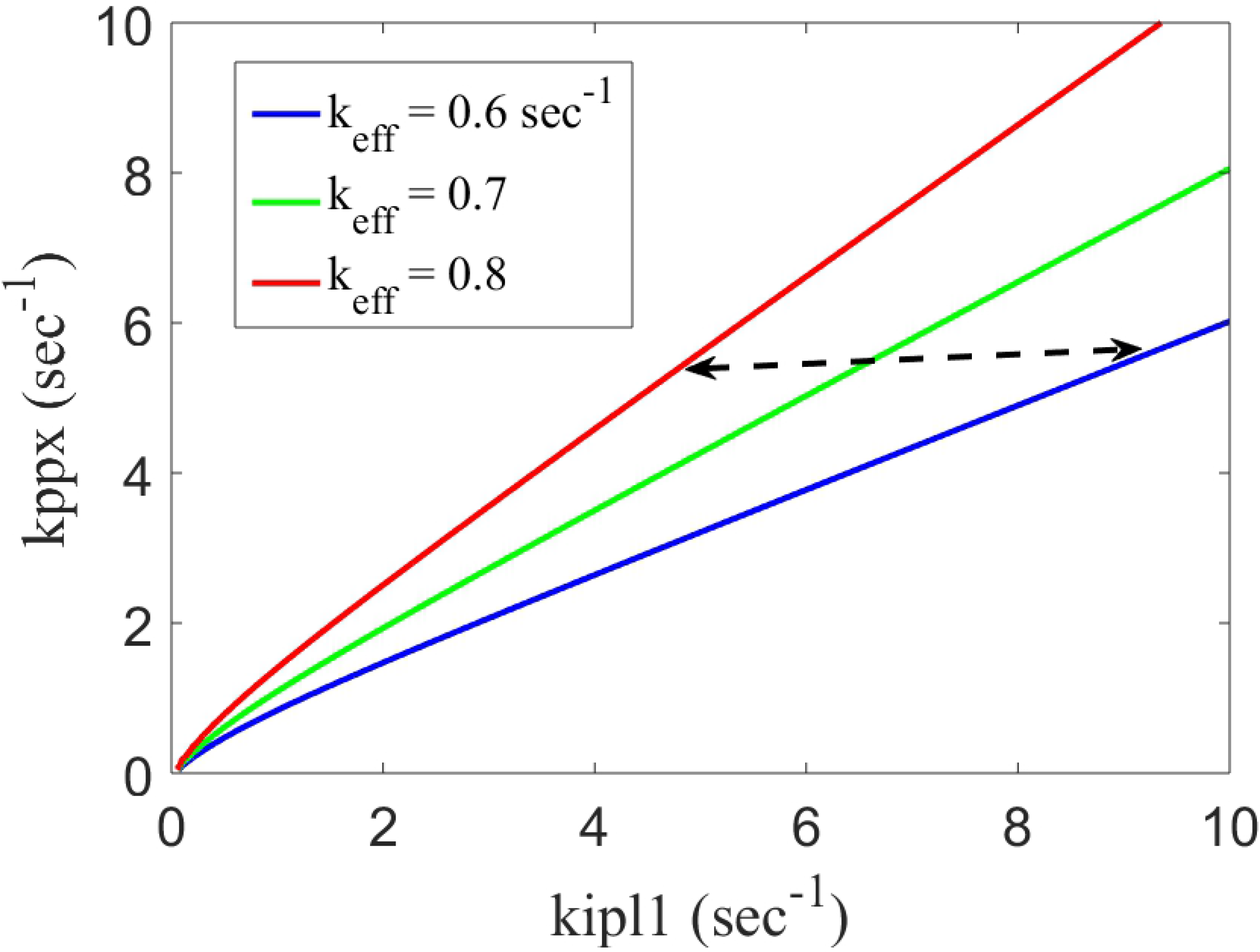

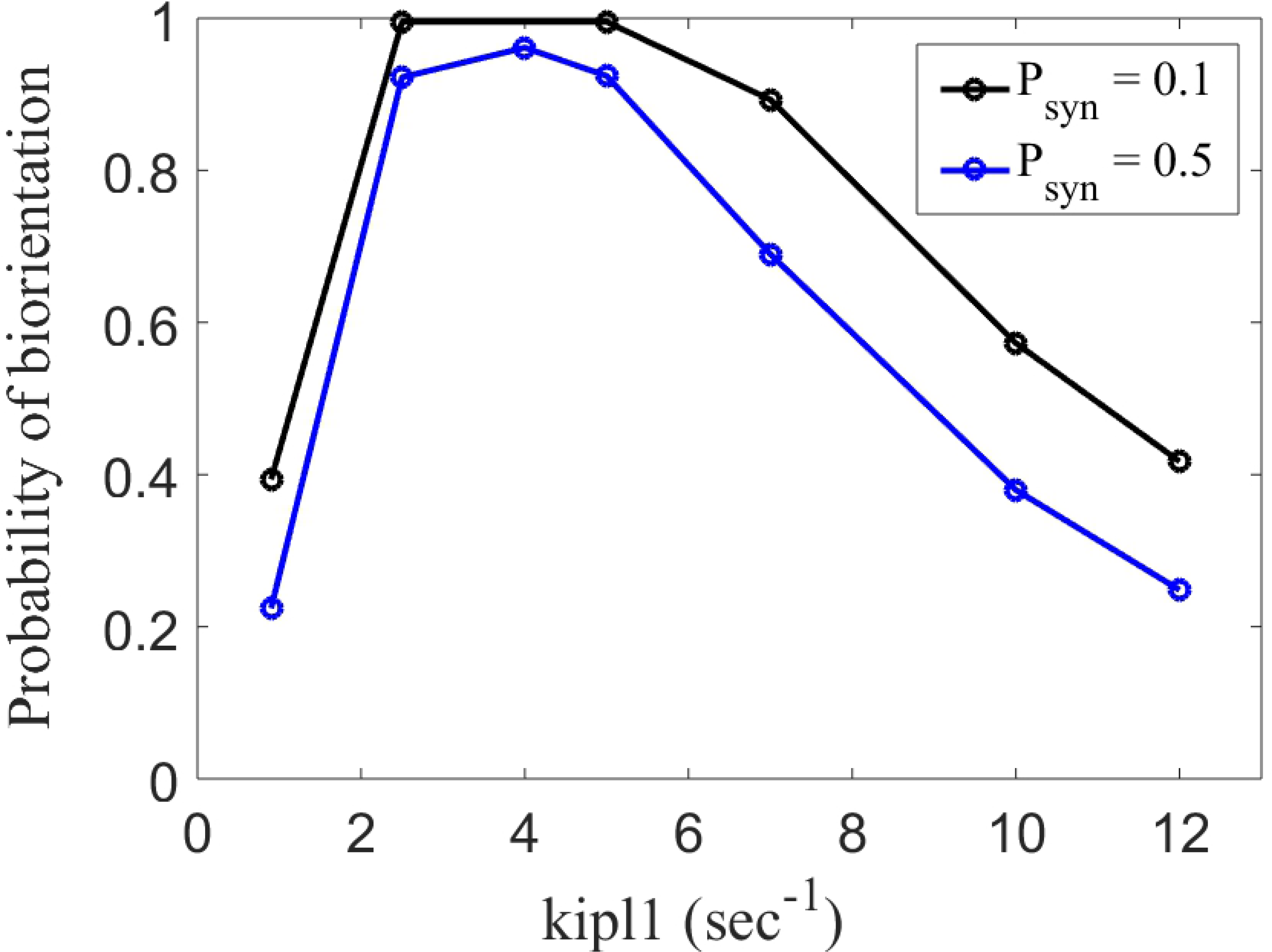

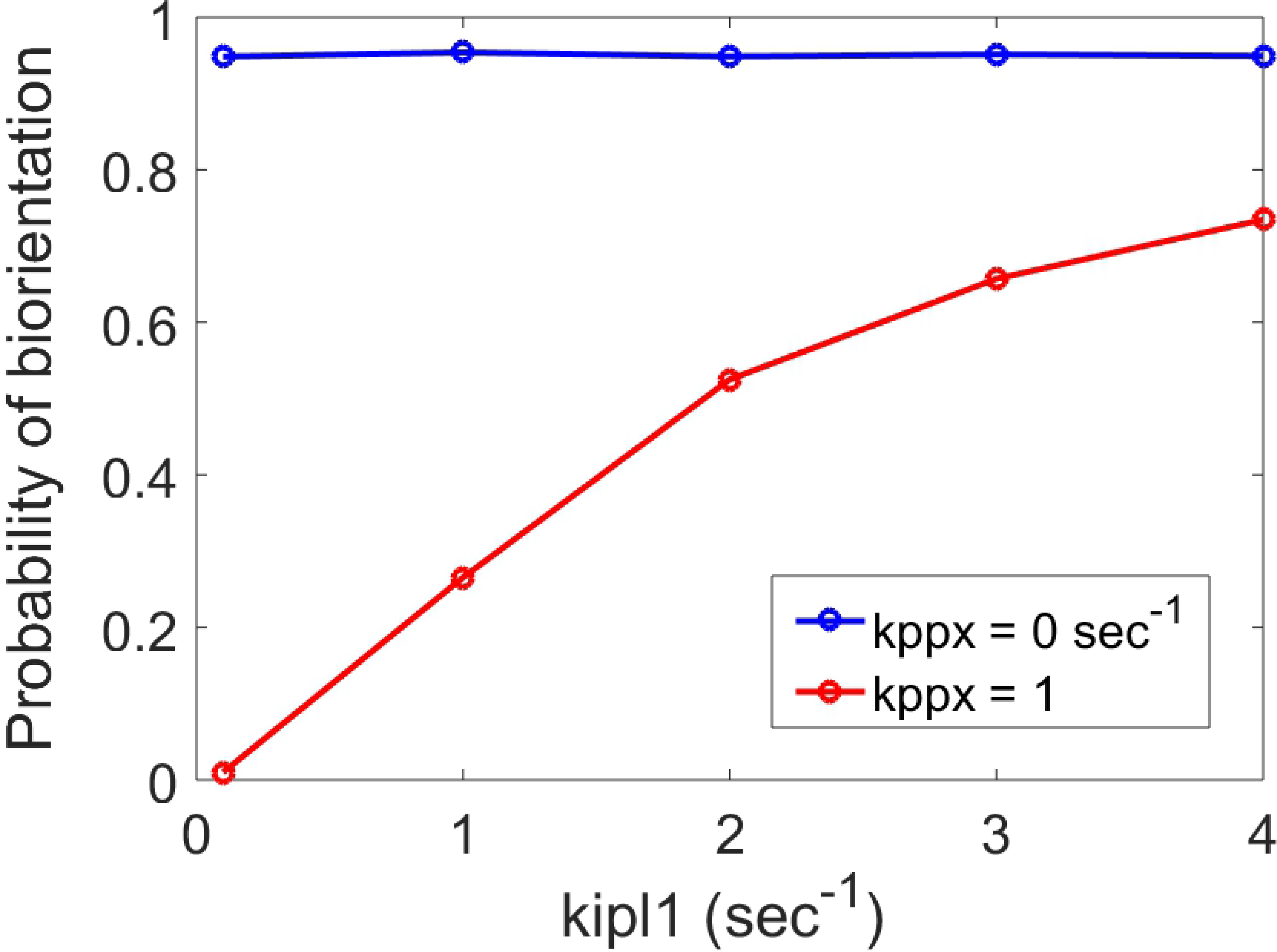

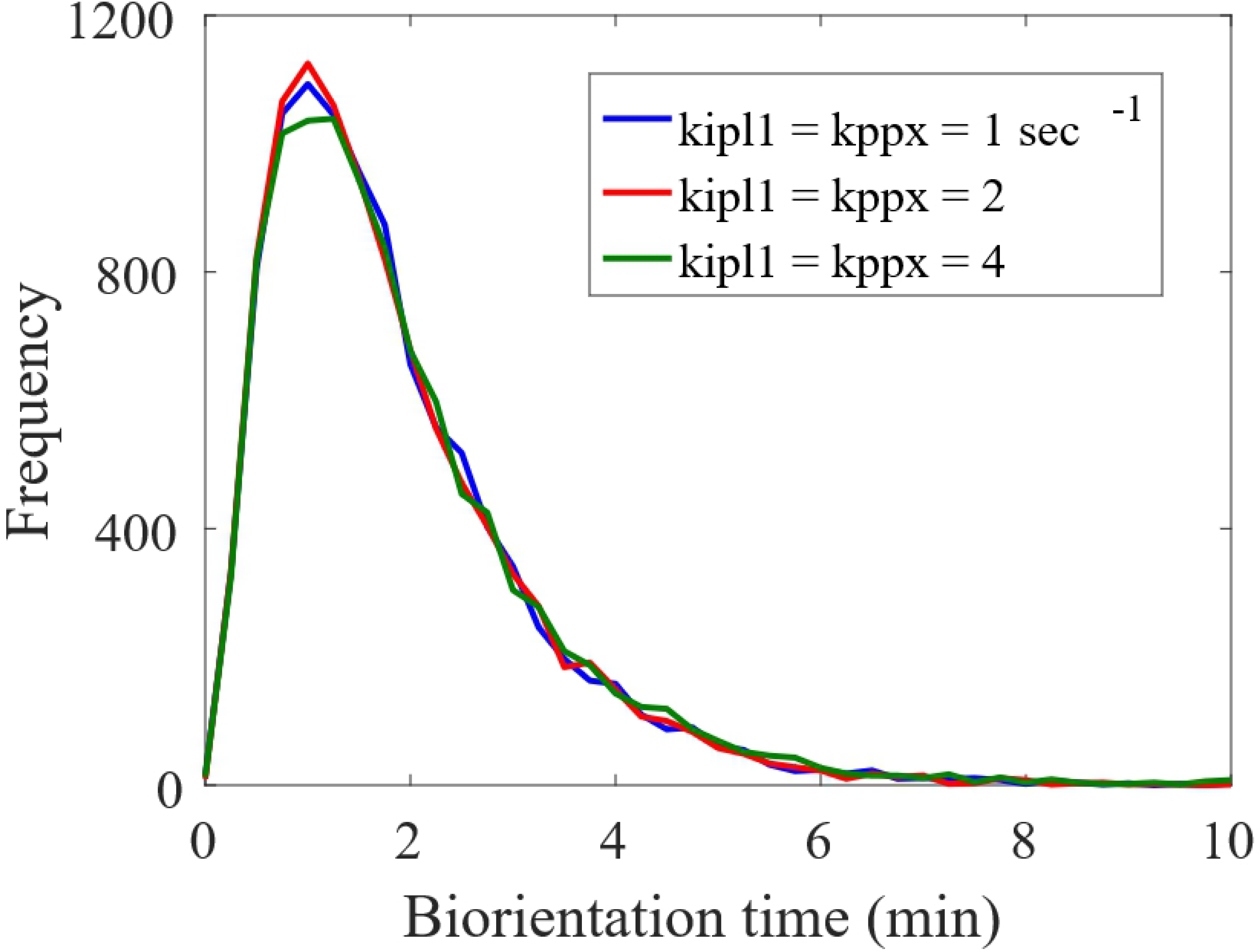

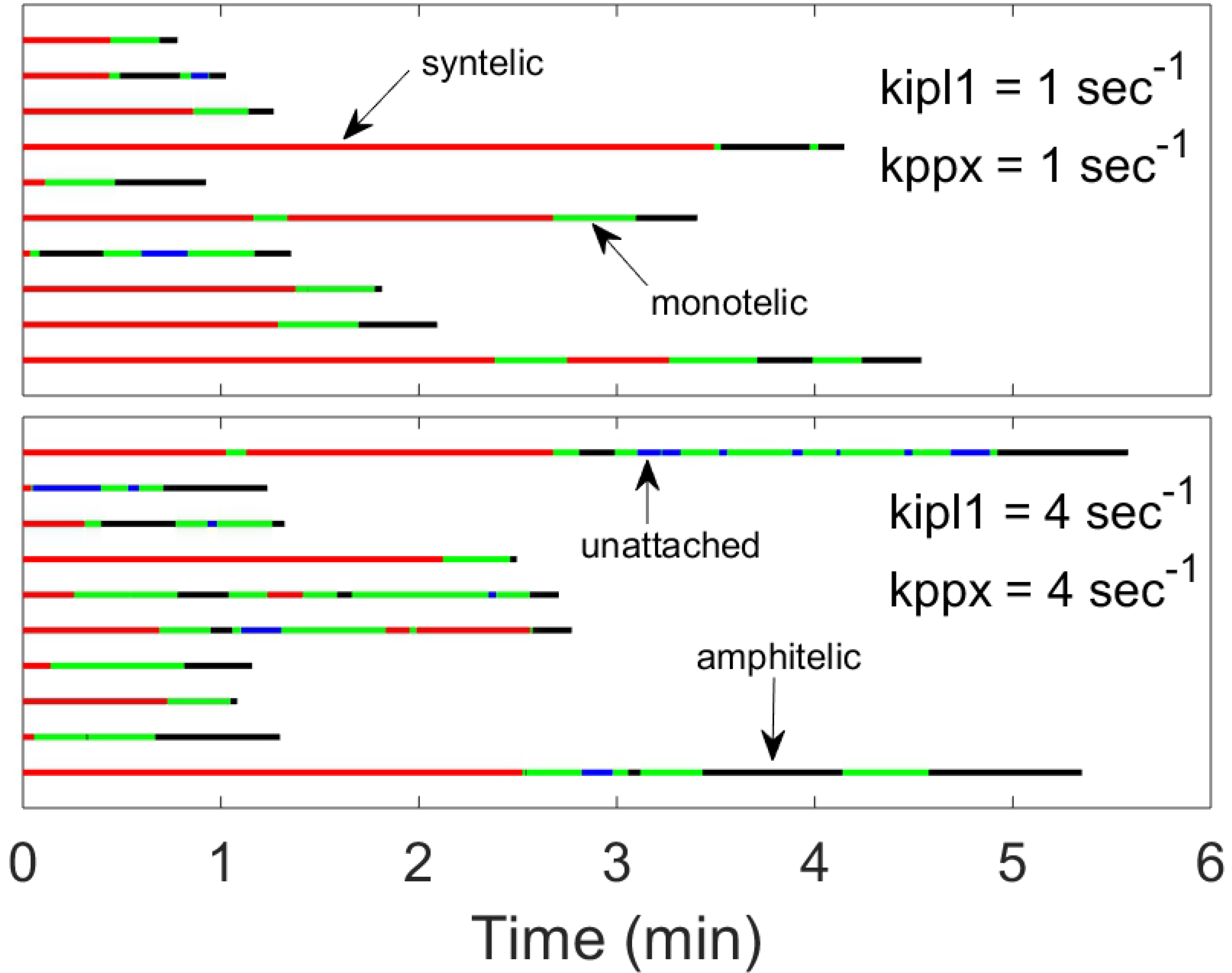

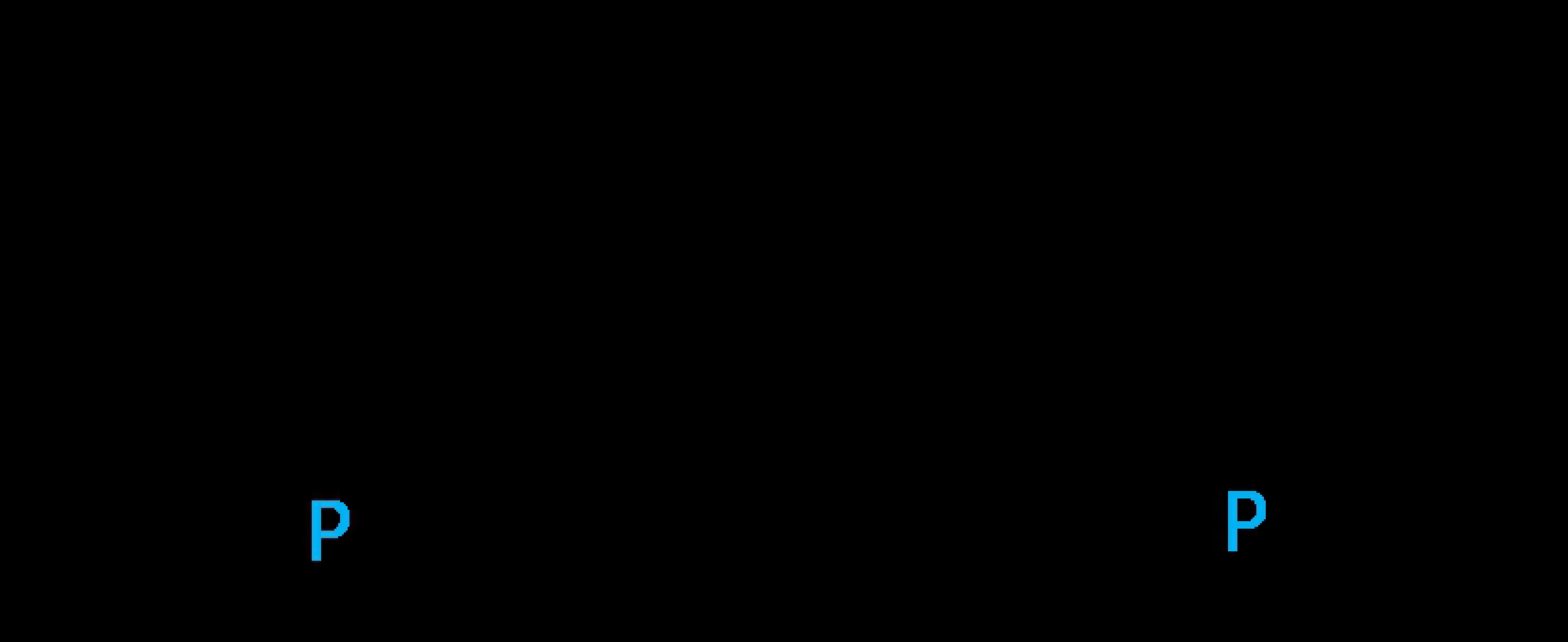

## Acknowledgements

“Research reported in this publication was supported by the National Institute of General Medical Sciences of the National Institutes of Health under Award Number R01GM078989. The content is solely the responsibility of the authors and does not necessarily represent the official views of the National Institutes of Health.

